# MultiFlow: coupled flow matching for predicting single-cell multiomic perturbation responses in unseen cellular contexts

**DOI:** 10.64898/2026.08.20.746112

**Authors:** Haochen Wang, Charming Zhang, Mengran Zhang, Xiaoming Nie, Qiao Liu

**Affiliations:** Department of Biostatistics, Yale University, New Haven, CT, USA; Stanford University School of Medicine, Stanford, CA; Program in Computational Biology and Bioinformatics, Yale University, New Haven, CT, USA

## Abstract

Predicting cellular responses to perturbation requires resolving coordinated changes across molecular layers, yet most single-cell perturbation models focus on transcriptional responses alone. Here we present MultiFlow, a coupled flow-matching framework that unifies generation and perturbation prediction of paired gene expression and chromatin accessibility. Through bidirectional information exchange between RNA and ATAC generative flows, MultiFlow learns how their joint distribution depends on cellular context and perturbation identity. Across multiomic generation benchmarks, MultiFlow accurately reproduced paired RNA-ATAC states and their population distributions. In multiome perturbation benchmarks, MultiFlow achieved the strongest overall performance in predicting both gene-expression and chromatin-accessibility perturbation responses, outperforming modality-specific perturbation-prediction methods in cell types whose perturbed states were withheld from training. Joint multiomic modeling further preserved perturbation-induced coordination between modalities, including concordant peak-gene effects and cross-modal cellular neighborhood structure. These results establish MultiFlow as a unified generative framework for modeling paired molecular states and predicting their coordinated perturbation responses across cellular contexts. Code and tutorial of MultiFlow are available at https://github.com/liuq-lab/MultiFlow.

## Introduction

Genetic perturbation provides a direct approach to interrogate gene function, uncover regulatory mechanisms and identify potential therapeutic targets ^1,2^. CRISPR-based single-cell perturbation technologies, such as Perturb-seq, extend this paradigm by resolving perturbation responses at single-cell resolution and revealing cellular heterogeneity that cannot be resolved by bulk measurements ^3,4^. However, perturbation responses are highly context dependent, with the same perturbation producing distinct molecular effects across cell types and cellular states ^5^. This creates a fundamental experimental challenge, as the combinatorial space of perturbations and cellular contexts far exceeds what can be practically measured experimentally. A central challenge is therefore to predict how a perturbation will reshape cellular states in contexts for which the response has not been experimentally observed ^6^.

A growing number of computational approaches have been developed to predict perturbation responses across cellular contexts, including variational autoencoder (VAE)-based methods ^7,8^, generative adversarial network (GAN)-based methods ^9^, optimal transport (OT) ^10^ and disentangled representation-learning frameworks ^11^. Recent benchmarking across diverse single-cell perturbation datasets has shown that trVAE ^7^, CellOT ^10^ and inVAE ^8^ are among the strongest-performing methods for cellular-context generalization, but also highlights substantial limitations in current approaches ^6^. Prediction performance declines when the target context becomes increasingly distinct from those observed during training, and current methods often emphasize average response prediction and may not fully capture population-level response distributions ^12^. Moreover, these methods characterize perturbation responses primarily through gene expression, ignoring the coordinated perturbation effect in other molecular layers, such as chromatin accessibility ^6,13^.

Genetic perturbations reshape cellular states through coordinated changes across molecular layers ^14^. Gene expression captures the transcriptional effect of perturbation, whereas chromatin accessibility reflects complementary changes in the regulatory state ^15^. Measuring both modalities within the same cell therefore provides a more holistic view of how perturbations reshape cellular programs. Recent experimental advances have made it possible to measure these responses jointly: Perturb-multiome ^14^ combines CRISPR-based perturbation with simultaneous profiling of chromatin accessibility and gene expression in the same cells, enabling perturbation-induced regulatory and transcriptional changes to be measured across cellular contexts. These advances create a pressing need for computational models that can predict perturbation responses jointly across molecular layers rather than treating each modality separately. This requires first learning the joint distribution of paired multiomic states ^16,17^ and then modeling how perturbations reshape each modality while preserving the cross-modal dependencies that reflect their underlying regulatory organization, particularly in unseen cellular contexts.

To address these challenges, we developed MultiFlow, a coupled flow-matching framework that unifies paired multiomic generation and perturbation-response prediction ^18^. Rather than modeling the two modalities independently, MultiFlow learns interacting RNA and ATAC flows that exchange information throughout the generative process, allowing perturbation-induced changes across molecular layers to be modeled as a coordinated response. By conditioning these coupled flows on cellular context and perturbation identity, MultiFlow learns context-dependent joint distributions of gene expression and chromatin accessibility. In the multiome generation benchmarks, MultiFlow reproduced the cell-type-specific joint distributions of gene expression and chromatin accessibility with high fidelity. In the multiome perturbation benchmarks, MultiFlow achieved the strongest overall prediction of both gene expression and chromatin accessibility responses compared to modality-specific methods, while preserving cross-modal relationships between RNA and ATAC modalities. Together, MultiFlow serves as a unified framework for generating paired multiomic states and predicting coordinated perturbation responses across cellular contexts.

## Results

### Overview of MultiFlow

MultiFlow is designed to jointly model paired multiomic states and their perturbation-induced transitions (**Fig. 1**). Modality-specific autoencoders are used to project each modality into a latent space. Starting from a shared Gaussian distribution where flow time (*t* = 0), MultiFlow learns two modality-specific flow trajectories that continuously transform noise toward paired RNA and ATAC states as flow time *tt* approaches one. This formulation preserves the distinct structure of each molecular modality while modeling them within a shared generative process. The key innovation is the bidirectional cross-flow module (CFM), which couples the RNA and ATAC trajectories throughout generation. Within each CFM, RNA and ATAC flows continuously exchange information through dual cross-attention ^19^, allowing the evolving state of each modality to inform the other. This design ensures the paired multiomic states are generated as a coordinated system rather than as independent molecular profiles.

**Figure 1.**
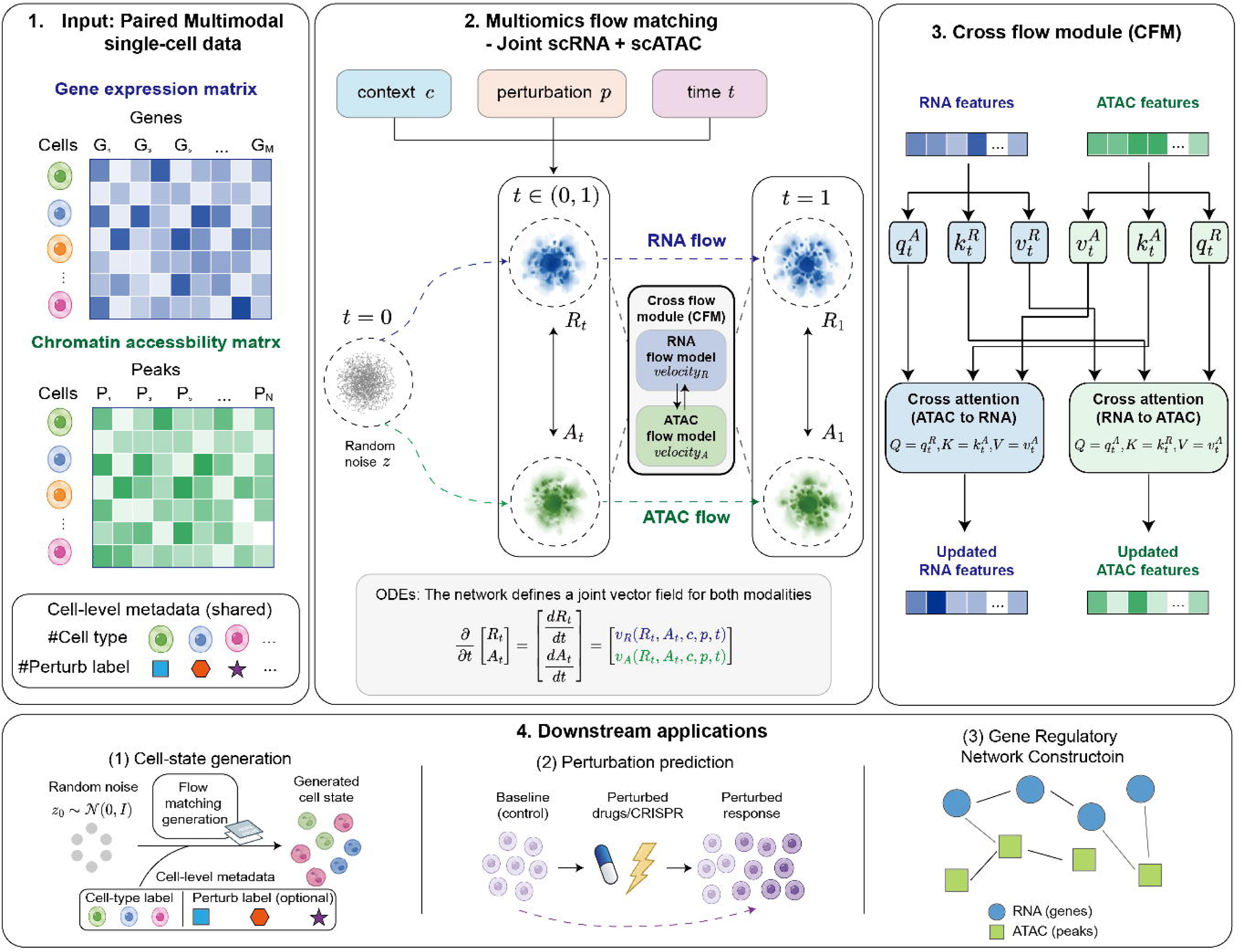
Overview of the MultiFlow framework. MultiFlow models paired single-cell multiomic profiles together with cellular context and perturbation information. RNA and ATAC profiles are projected into modality-specific latent representations and generated through two coupled flow-matching trajectories that evolve from a shared Gaussian reference distribution toward paired multiomic cell states. The RNA and ATAC flows are connected by bidirectional Cross Flow Modules (CFMs), which use dual cross-attention to exchange information between modalities throughout the generative process while preserving modality-specific dynamics. The learned cross-modal representations can further be analyzed to characterize regulatory relationships in new cellular contexts.

MultiFlow unifies multiomic generation and perturbation-response prediction within a unified generative framework. For multiomic cell-state generation, the coupled flows learn cell-type-specific paired RNA-ATAC joint distributions under the unperturbed condition. For perturbation prediction, the same flow-matching system is built conditioned on perturbation and a control-derived cellular-state representation, enabling perturbation effects learned across observed cellular contexts to be transferred to unseen cellular contexts. Through both multiomic generation and perturbation benchmarks, we demonstrate that MultiFlow could faithfully reproduce the joint distribution of paired RNA-ATAC states and accurately predict coordinated gene expression and chromatin accessibility perturbation responses in unseen cellular contexts, compared with competing methods. Together, these results establish MultiFlow as a unified generative framework for learning paired multiomic states and predicting how they are jointly reshaped by perturbation across cellular contexts.

### MultiFlow captures cell-type-specific paired multiomic distributions

We first evaluated whether MultiFlow could faithfully generate cell-type-specific RNA and ATAC states before evaluating its ability to predict perturbation responses. In this setting, cell-type identity specifies the cellular context, while the perturbation condition is kept inactive, making cell-state generation a special case of the same generative framework under a null perturbation. The coupled flows therefore learn the distribution of unperturbed paired multiomic states for each cell type.

We benchmarked MultiFlow against the state-of-the-art multiomic generative models, including scDiffusion-X ^20^, CFGen ^21^, and MultiVI ^22^, using the OpenProblems multiome dataset ^23^. We additionally compared MultiFlow with scDesign3 ^24^, a statistical simulator for single-cell multiomics, on the smaller PBMC10k dataset ^25^, due to its limited computational scalability on large-scale datasets. For each dataset, 80% of cells were used for training and the remaining 20% were held out for evaluation. We evaluated complementary aspects of generation fidelity, including preservation of cell-state organization, agreement between real and generated population distributions, maintenance of conditioned cell-type identity, and feature-level agreement of cell-type-averaged RNA and ATAC profiles (see Methods).

On the OpenProblems dataset, cells generated by MultiFlow closely matched the cell-type organization of the held-out data and overlapped extensively with real cells in the shared UMAP ^26^ representation (**Fig. 2a**). We quantified the agreement between real and generated populations using complementary mean-profile and distribution-level metrics. For RNA, MultiFlow achieved the highest iLISI of 0.898, outperforming the best competing method scDiffusion-X by 0.052 (**Fig. 2b** and **Supplementary Fig. 1**). The advantage was even stronger for chromatin accessibility, for which MultiFlow achieved both the highest iLISI ^27,28^ (0.917) and the lowest MMD ^29^ (0.011). Together, these results show that MultiFlow accurately captures population-level structure of multiomic profiles.

**Figure 2.**
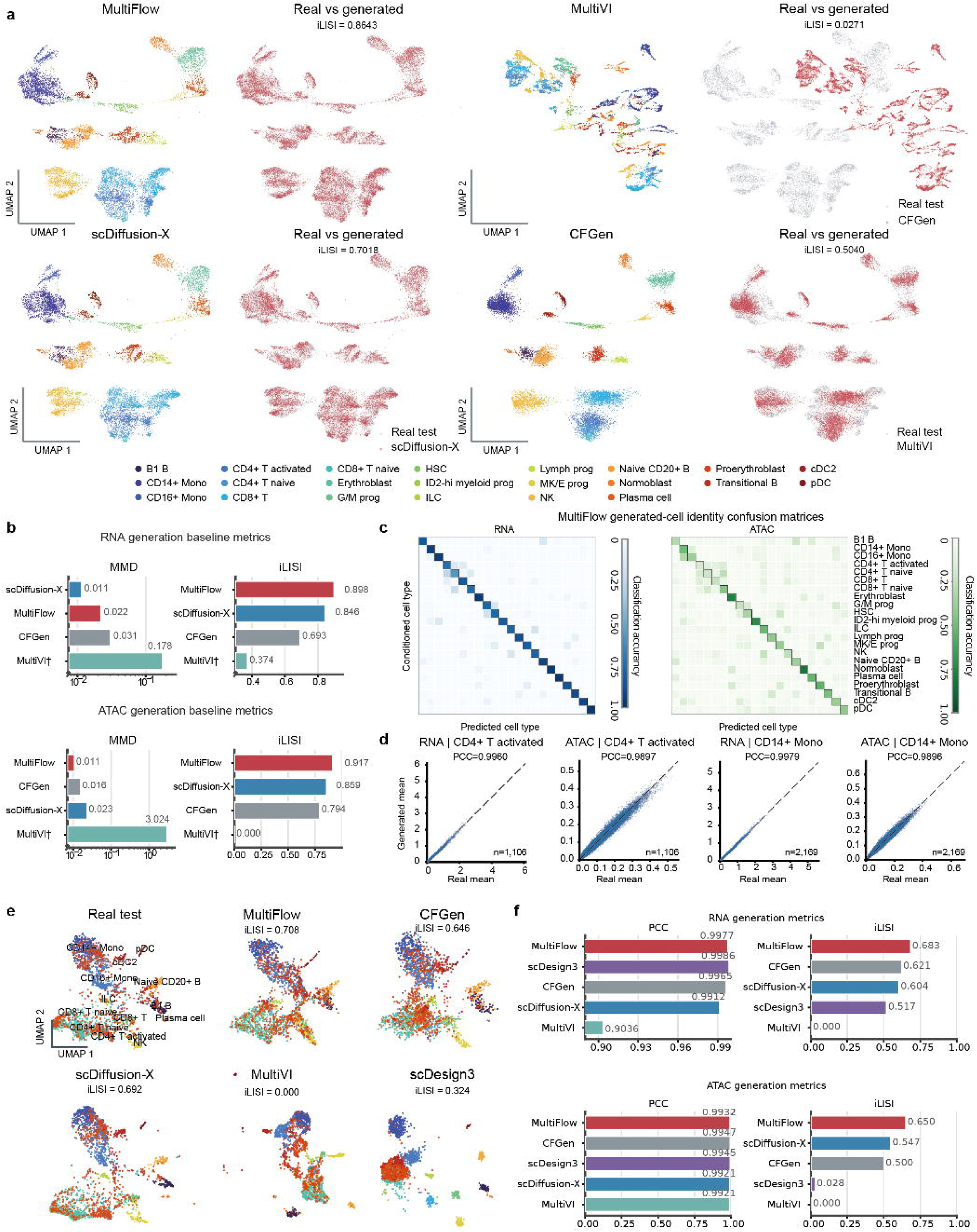
MultiFlow generates cell-type-specific paired multiomic states. **a,** UMAP visualization of held-out real and generated multiomic profiles by different methods using the OpenProblems dataset. Cells are colored by cell type, while overlay panels only distinguish real and generated cells. **b**, Distribution-level agreement between real and generated RNA and ATAC profiles measured by maximum mean discrepancy (MMD) and integration local inverse Simpson’s index (iLISI). **c**, RNA and ATAC cell-type identity confusion matrices for MultiFlow-generated cells. Rows represent conditioned cell types and columns represent identities predicted by external classifiers trained on real cells. **d**, Scatter plots between real and generated cell-type-averaged molecular profiles for two representative cell types. Each point represents a gene or peak. Pearson correlation coefficient (PCC) was calculated for each panel. **e**, UMAP visualization of held-out real and generated cells by different methods using the independent PBMC10k dataset. Cells are colored by cell type. **f**, PCC and iLISI comparisons on PBMC10k dataset for RNA and ATAC generation, respectively.

Next, we examined whether the generated profiles retained the molecular identity of the cell type. To quantify cell-type fidelity, we trained cell-type classifiers on real multiome profiles and applied them to the corresponding generated cells. The resulting confusion matrices showed high classification accuracy along the diagonal for both modalities, indicating that MultiFlow preserved cell-type-specific patterns (**Fig. 2c**). MultiFlow also closely recovered cell-type-averaged molecular profiles across different cell types (**Fig. 2d** and **Supplementary Fig. 2**). For example, Pearson correlations between real and generated feature means reached 0.9960 for RNA and 0.9897 for ATAC in CD4+ T cell type. Additional paired cross-modal analyses are provided in **Supplementary Fig. 3**, which showed that MultiFlow captures paired RNA-ATAC coupling while preserving the marginal distribution of each modality.

We further tested whether these properties generalized to an independent multiome dataset. On PBMC10k, MultiFlow reproduced the major immune-cell populations and showed close agreement with the held-out cells in UMAP ^26^ visualizations (**Fig. 2e**). All leading methods achieved high mean-profile correlations but more diverse performance in distribution-level metrics (**Fig. 2f** and **Supplementary Fig. 4**). For instance, MultiFlow achieved the highest iLISI for both ATAC (0.650) and RNA (0.683), leading to a substantially relative 18.83% improvement in ATAC over the best baseline scDiffusion-X. A similar improvement was also observed in RNA modality. These results indicate that accurate recovery of population-average molecular profiles does not necessarily translate into faithful recovery of single-cell distributions. Collectively, the multiomic generation benchmarks establish the ability of MultiFlow to generate coherent cell-type-specific multiome profiles, providing the generative foundation for predicting how paired multiomic states respond to perturbation.

### MultiFlow predicts multiomic perturbation responses in unseen cellular contexts

After establishing that MultiFlow can generate unperturbed faithful multiomic profiles, we next asked whether the same coupled flow-matching generative framework could predict how these states are reshaped by perturbation in unseen cellular contexts. In this setting, the perturbation condition is activated, while the cellular context is represented by the unperturbed control cells. This allows MultiFlow to transfer perturbation responses learned from observed cellular contexts to unseen target contexts. We evaluated this capability using the Perturb-multiome dataset in a leave-one-cell-type-out setting, in which all perturbed cells from one cell type were excluded from training. We evaluated prediction at three levels: transcriptional response, chromatin-accessibility response and the coordination between the two molecular layers.

### MultiFlow improves RNA perturbation prediction

We first examined transcriptional response prediction, for which a broad range of single-cell perturbation methods are available for comparison. Across held-out cell types, MultiFlow achieved the best RNA perturbation prediction performance across all four evaluated criteria, including perturbation-effect and population-distribution metrics (**Fig. 3a** and **Supplementary Fig. 5,6**). On the top-100 perturbation-responsive genes, MultiFlow substantially increased iLISI from 0.716 for the strongest baseline to 0.867, a 21.1% relative improvement, and reduced the MMD 2.7-fold from 0.093 to 0.034 over the strongest baseline. These improvements were further supported by UMAP visualizations of a specific perturbation where MultiFlow predictions closely followed the observed transcriptional response across cell types while several competing methods showed greater distortion (**Fig. 3c** and **Supplementary Fig. 7**).

**Figure 3.**
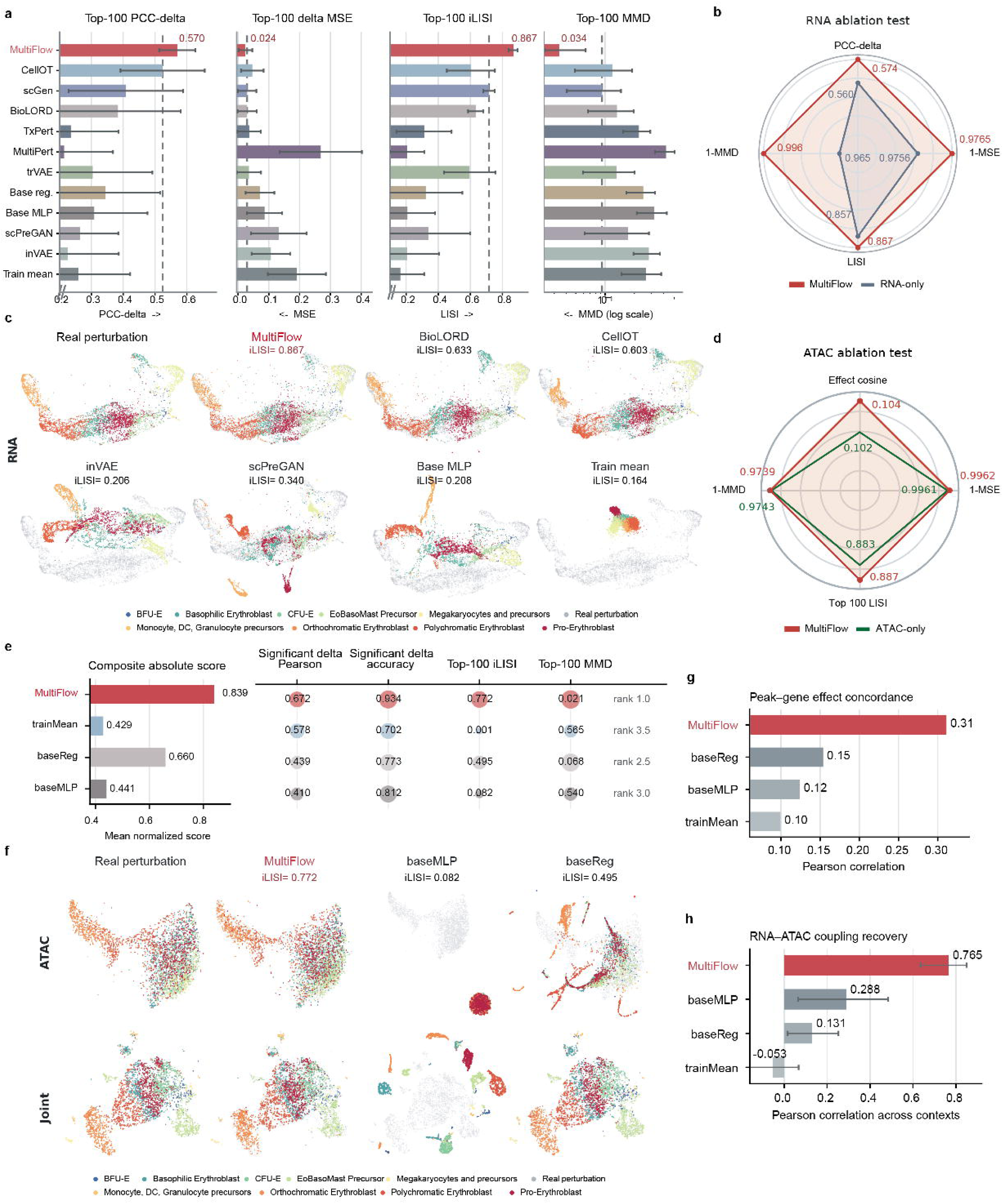
MultiFlow predicts coordinated RNA and ATAC perturbation responses in unseen cellular contexts. **a**, Leave-one-cell-type-out benchmark of RNA perturbation prediction using PCC-delta, delta MSE, iLISI, and MMD on the top-100 perturbation-responsive genes. Bars show means across held-out cell types, and error bars indicate between-cell-type variation. **b**, RNA perturbation prediction comparing the full MultiFlow model with an RNA-only model, assessing the contribution of chromatin accessibility information to transcriptional response prediction. Metrics are oriented so that greater radial extent indicates better performance in the radar plot. **c**, Shared UMAP visualization of observed and predicted RNA profiles under the FOSL1 perturbation across held-out cell types. **d**, ATAC perturbation prediction comparing the full MultiFlow model with an ATAC-only model, assessing the contribution of RNA information to chromatin accessibility response prediction. Metrics are oriented so that greater radial extent indicates better performance in the radar plot. **e**, Evaluation of predicted ATAC responses using significant-delta correlation and directional accuracy, Top-100 iLISI and MMD, and their composite absolute score. **f**, UMAP visualization of observed and predicted profiles under the FOSL1 perturbation across held-out cell types in ATAC space and a joint RNA-ATAC space for MultiFlow and baselines. **g**, Peak-gene effect concordance, measuring the agreement between predicted chromatin accessibility changes and transcriptional responses of nearby genes. **h**, Recovery of cell-level RNA-ATAC coordination, measured by the correlation between real and predicted context-level mean Jaccard overlap of RNA- and ATAC-defined nearest-neighbor sets.

We next asked whether modeling chromatin accessibility jointly with gene expression contributed to transcriptional response prediction. Removing the ATAC flow produced a consistent reduction in RNA prediction performance across nearly all evaluated metrics (**Fig. 3b**), indicating that chromatin accessibility provides additional information beyond the transcriptomic state for transferring perturbation responses across cellular contexts.

### MultiFlow predicts chromatin accessibility responses in unseen cellular contexts

We then assessed whether MultiFlow could recover the chromatin accessibility response. MultiFlow ranked first overall and achieved a composite absolute score of 0.839, compared with 0.660 for the best baseline (**Fig. 3e** and **Supplementary Fig. 8**). MultiFlow recovered both the magnitude and direction of accessibility changes, achieving a correlation of 0.672 for perturbation-induced changes at significant peaks and a directional accuracy of 0.934. It also captured the single-cell distribution of chromatin responses with a top-100 iLISI of 0.772 and an MMD of 0.021, all surpassing the best baseline by a large margin. Consistent with these quantitative results, ATAC and joint RNA-ATAC UMAP visualizations for a specific perturbation showed that MultiFlow predictions closely followed the observed perturbed cell profiles, whereas baseline predictions showed substantially greater deviations (**Fig. 3f**).

We further examined whether coupling to the RNA modality contributed to ATAC response prediction. Compared with an ATAC-only model, the full MultiFlow model showed modest improvements in effect cosine similarity, MSE and iLISI, although MMD was slightly worse (**Fig. 3d**). Notably, these gains were smaller than the improvements observed when adding ATAC information to RNA perturbation prediction, suggesting an asymmetric contribution of cross-modal information.

Together with the RNA results, these findings show that MultiFlow can transfer perturbation effects across cellular contexts at both transcriptional and chromatin accessibility levels.

### MultiFlow preserves perturbation-induced RNA-ATAC coordination

Accurate prediction of the two modalities separately does not necessarily ensure that their cross-modal relationship is preserved in the predicted multiomic state. We therefore examined whether perturbation-induced RNA-ATAC relationships are preserved at both genomic-feature and cellular-neighborhood levels. First, we asked whether predicted accessibility changes were spatially concordant with transcriptional responses of nearby genes. MultiFlow achieved a peak-gene neighborhood-effect score of 0.31, which is more than twice higher than the strongest baseline (0.15), indicating substantially stronger agreement between predicted chromatin changes and nearby transcriptional effects (**Fig. 3g**). We next assessed cross-modal organization at the single-cell level by comparing nearest-neighbor structures independently defined from RNA and ATAC profiles. The correlation between real and predicted RNA-ATAC neighborhood-coupling scores reached 0.765 for MultiFlow, compared with 0.288 for the strongest baseline, whereas several baselines showed little or negative correspondence (**Fig. 3h**). These complementary analyses indicate that MultiFlow preserves perturbation-induced relationships between regulatory and transcriptional states rather than just capturing the marginal responses of RNA and ATAC separately.

Collectively, these results show that MultiFlow generalizes perturbation responses to unseen cellular contexts at three interconnected levels: transcriptional changes, chromatin accessibility changes and the cross-modal relationships. By predicting how a paired RNA-ATAC state is jointly reshaped under a specific perturbation, MultiFlow extends single-cell perturbation prediction from a transcriptome-centered task to coordinated multiomic response modeling.

### MultiFlow recovers peak-gene associations and functional gene modules

Having established that MultiFlow preserves coordinated RNA-ATAC perturbation responses, we next asked whether the model captures interpretable regulatory associations during its cross-modal computation. In MultiFlow, the RNA and ATAC flows exchange information through the dual cross-attention of the cross-flow modules (CFMs). The attention maps offer a direct view of how the two modalities are connected and interacted with each other. To interpret what the attention map learned, we used a gradient-based strategy to trace its hidden elements back to peaks and genes (Methods). The cross-attention map from the CFM connects 512 ATAC and 512 RNA hidden elements, and each entry of the cross-attention map links one ATAC element to one RNA element. For each hidden element, the gradient with respect to an input feature (gene or peak) of its own modality measures how strongly the element responds to a change in that feature, so ranking the features by this gradient assigns each ATAC element with its top-attributed peaks and each RNA element with its top-attributed genes, and thereby defines candidate peak-gene association links (**Fig. 4a**).

**Figure 4.**
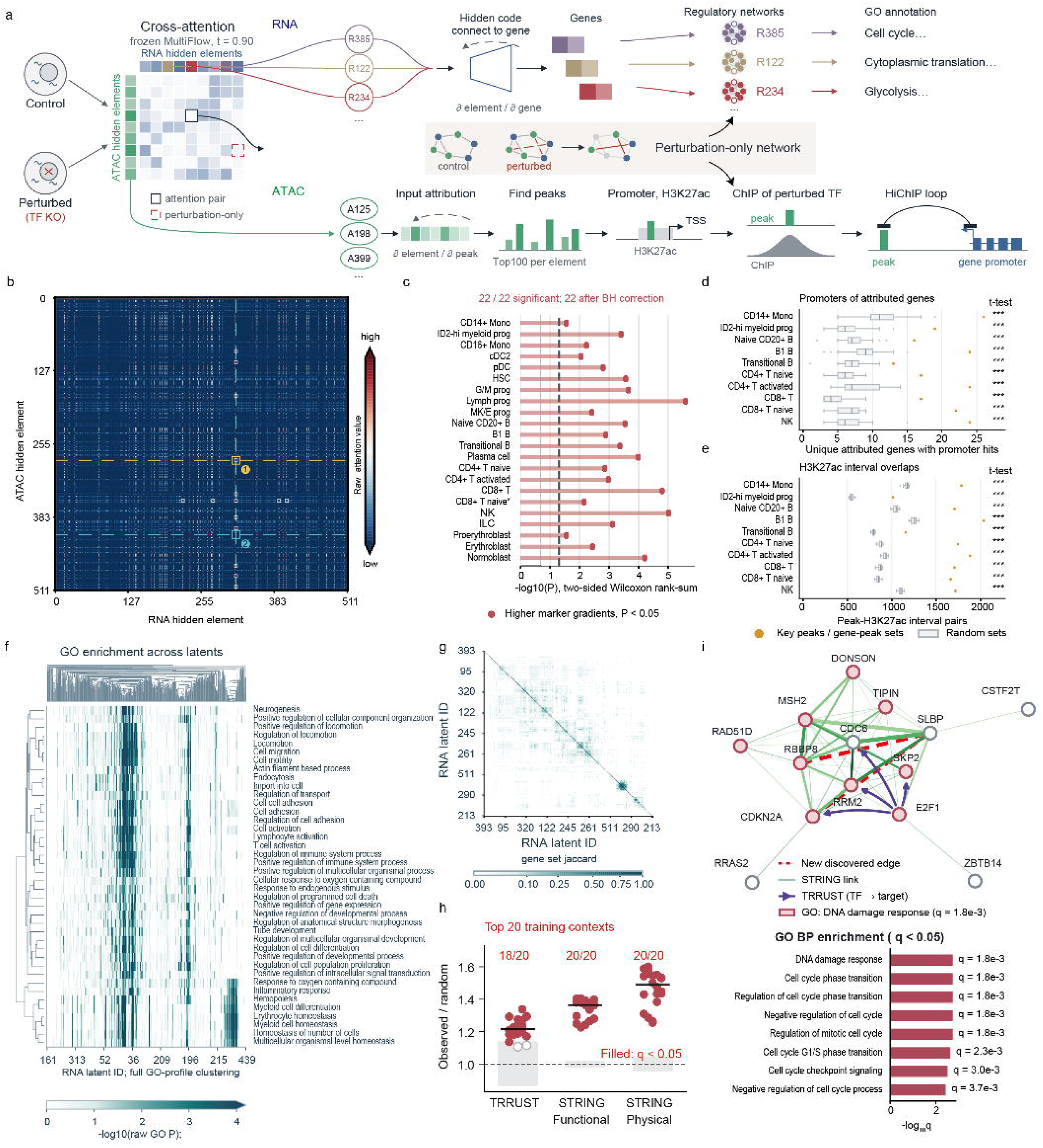
MultiFlow’s internal attention, attribution and hidden code recovery. **a**, Schematic of the analysis: cross-attention between the ATAC and RNA hidden elements of the trained model in the multiome or multiomic perturbation datasets. **b**, Attention map of CD14⁺ monocytes; white squares mark the 15 strongest pairs, and numbers 1 and 2 in the figure show the strongest pairs among the top 0.1% that link the *IL1B* and *S100A9* promoters to these genes. **c**, Attribution of marker genes versus all other genes (13,431 measured genes) in 22 cell types, scored by the mean absolute gradient over the 20 most-attended RNA elements; x axis, −log10(P) of a two-sided Wilcoxon rank-sum test; dashed line, *p* = 0.05. **d**, Attributed genes whose promoter overlaps an attributed peak in ten cell types (gold dots), compared with 20 size-matched random sets (box plots). **e**, Overlaps between attributed peaks and H3K27ac ChIP-seq intervals in the same cell types. Asterisks show a one-sided t-test against the 20 random sets. **f**, GO enrichment of RNA hidden elements in the *GATA1* perturbation in Baso-E cell type. **g**, Gene-set similarity of RNA hidden elements in the *GATA1* perturbation in Baso-E cell type. **h**, Enrichment of TRRUST and STRING pairs among genes co-selected by an RNA element in 20 training contexts. **i**, Gene module of R399 RNA latent variable; red dashed lines are new candidate edges. The module contains more genes connected by STRING (score ≥ 400) or TRRUST than expected by chance (14 versus 3.8; *p* = 0.003) and more TRRUST pairs (4 versus 0.06; *p* = 0.001), and *p* values are from a one-sided permutation test that shuffles gene names.

We first applied this interpretation analysis to the cell-state generation model trained on the OpenProblems multiome dataset and asked whether these links agree with known regulatory biology. **Fig. 4b** shows an example of the attention map, averaged over the CD14⁺ monocytes of the OpenProblems multiome data. We chose this cell type because its marker genes are well characterized ^23^ and a matched CD14⁺ monocyte H3K27ac ChIP-seq profile is available ^20^. Among the top 0.1% of the attention pairs averaged over CD14^+^ monocytes, we show, for each of two classical monocyte genes, *IL1B* and *S100A9*, the strongest pair that links a peak at its promoter to the gene itself. In pair A284-R316, which ranks seventh (top 0.003%), the top-attributed peaks of ATAC element A284 were located in the *IL1B* promoter region, and the attributed genes of RNA element R316 included *IL1B*. Another pair, A414-R316, which ranks 31st (top 0.01%), linked the *S100A9* promoter to *S100A9* in the same way. For both genes, the attributed ATAC peak spanned the corresponding TSS and overlapped an H3K27ac-enriched region in CD14⁺ monocytes ^30^ (**Supplementary Fig. 9**), so the model connected the accessibility of each gene’s active promoter to the gene’s expression.

We next asked whether attribution captured cell identity beyond individual examples. For each of the 22 cell types, we compared the attribution of its marker genes (2 to 12 per cell type) with that of all other genes, and we found that marker genes carried higher attribution than all other genes in all 22 cell types after Benjamini-Hochberg (BH) correction (**Fig. 4c**, Methods). Although ATAC and RNA attribution are computed separately, in all ten cell types with a matching H3K27ac ChIP-seq profile, the attributed peaks fell at the promoter regions of attributed genes 2.1- to 3.4-fold more often than in size-matched random peak sets (median 3.0-fold; **Fig. 4d**) and overlapped H3K27ac-marked enhancer regions 1.5- to 2.0-fold more often (median 1.7-fold; **Fig. 4e**). Both enrichments were significant in all ten cell types against 1,000 random sets after BH correction (Methods). The chromatin accessibility and gene expression features that the model relies on therefore converge on the same active promoters.

Assembling these links further recovered monocyte biology. In CD14⁺ monocytes, we used the top 0.1% of attention pairs to link attributed peaks and genes. We retained links only when the peak overlapped both the corresponding gene’s promoter and an H3K27ac-enriched region. This yielded 245 paths through 152 pairs that contain 71 genes (**Supplementary Fig. 10a**). These genes were enriched for antigen presentation and immune effector processes (248 GO terms at q < 0.05; for example, regulation of immune effector process, q = 6 × 10⁻⁴; **Supplementary Fig. 10b**), consistent with the antigen-presenting and inflammatory functions of monocytes ^31,32^.

We next applied the same analysis to the perturbation-prediction model trained on the Perturb-multiome dataset ^14^. We used the trained model to generate cells of each training context in the perturbation state. For each RNA element, attribution identified the genes that the element links, which form its gene set (Methods). These gene sets mapped to distinct Gene Ontology programs ^33^. For example, in *GATA1*-perturbed basophilic erythroblasts (Baso-E), the enriched GO terms included cell cycle (R385), cytoplasmic translation (R122) and glycolysis (R234) (**Fig. 4f** and **Supplementary Figs 11-12**). Gene sets associated with different RNA elements showed little overlap overall (only 1.6% of pairs with Jaccard index ≥ 0.1), although small clusters of elements shared subsets of genes (**Fig. 4g**). Comparison of GO enrichment profiles further revealed functional similarities among elements despite limited gene-set overlap (median cosine similarity: 0.21; **Supplementary Fig. 13**). These findings suggest that related biological functions are represented across multiple RNA elements with largely distinct gene sets.

We then asked whether genes grouped by the same element were connected by known regulatory and functional relationships. For each perturbation, cell-type combination, we linked genes assigned to the same RNA element and weighted each edge by the number of elements selecting both genes (Methods). Across the 20 training perturbation and cell-type combinations in which the model predicted the perturbation effect best (highest RNA PCC-delta), gene pairs selected by the hidden element were enriched for known transcription factor (TF)-target pairs in TRRUST ^34^ (18/20 combinations) and for functional and physical protein associations in STRING ^35^ (20/20 for each), relative to gene-label permutations that preserved gene-selection frequencies (BH-adjusted P < 0.05; **Fig. 4h**). Similar enrichment patterns were observed when the analysis was restricted to the top 5, 10 or 15 combinations (**Supplementary Fig. 14**). At the level of individual RNA elements, selected modules associated with pyruvate metabolism, cell division, erythrocyte development and DNA replication also contained more reference-supported gene pairs (**Supplementary Figs. 15-16**).

Finally, we examined one module in detail. In the RNA element R399 of *SPI1*-perturbed polychromatic erythroblasts (Poly-E), 14 genes were connected by STRING or TRRUST, including four TRRUST targets of E2F1 (*CDC6*, *CDKN2A*, *RRM2* and *SKP2*), and the module was enriched for DNA-damage-response and cell-cycle terms (**Fig. 4i**). Within this module, there are 18 gene pairs co-selected by at least three RNA elements (Methods). 16 out of 18 (88.9%) gene pairs had a STRING or TRRUST support. The other two, *CDKN2A-SLBP* and *RBBP8-SLBP* lacked support in existing databases and were retained as new candidate associations. These two pairs connect genes involved in replication-dependent histone mRNA metabolism (SLBP ^36^), cell-cycle control (CDKN2A ^37^) and the DNA end resection (RBBP8 ^38^). Given the coupling relationship between histone mRNA metabolism and DNA replication ^36^, we interpret these recurrent links as potential evidence of functional coordination between histone production, DNA repair and cell-cycle control.

Together, these results indicate that MultiFlow’s internal representations are not only predictive but also interpretable. In the multiome dataset, attention and attribution recover marker programs, promoter and H3K27ac support and an attention-derived geme regulatory network. In the Perturb-multiome analysis, the hidden elements form functional modules whose gene pairs are supported by curated interactions, and these modules also propose new candidate links that current databases do not yet contain.

### TF knockouts re-weight attention toward target genes and TF-binding sites

We next asked which parts of the attention-derived network were specific to perturbation. For each training perturbation and cell-type combination, we applied the trained model to real non-targeting (NT) and knockout cells to construct control and perturbation networks. We then compared the two networks and isolated the perturbation-only subnetworks for functional analysis and validation against TF-binding data (**Fig. 5a**).

**Figure 5.**
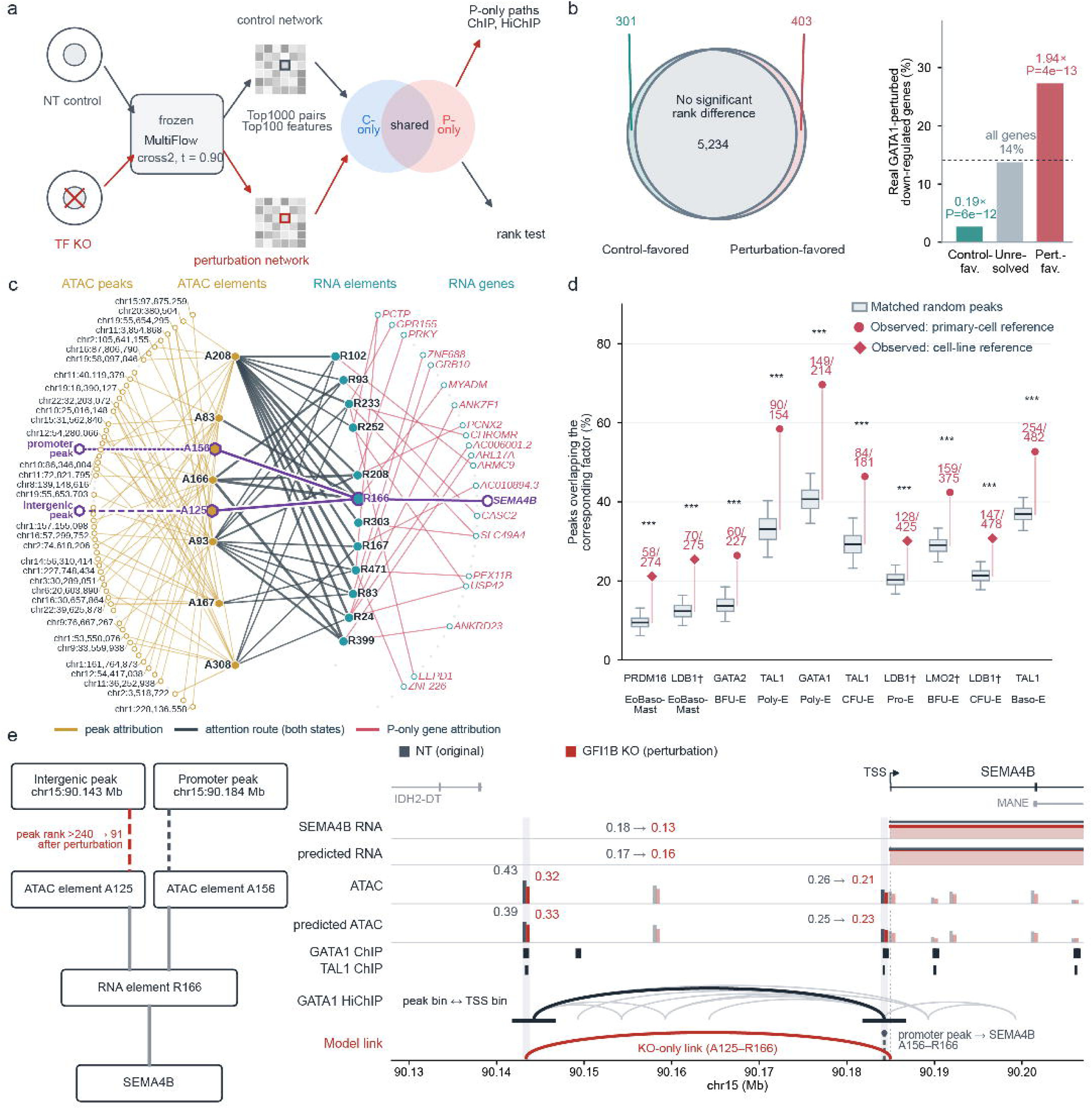
TF knockouts shift MultiFlow’s attention network toward knockout-responsive genes and factor-bound peaks. a, Schematic of the analysis: control and knockout cells of a training context are passed through the trained model (CFM2, t = 0.90; Top1000 attention pairs and Top100 attributed features per element). The control and perturbation networks are compared to find routes. **b**, Genes classified by the change in their attribution rank after *GATA1* knockout in Baso-E (left), and the share of each class that is down-regulated in observed knockout cells (right; dotted line, all tested genes). **c**, Perturbation-only gene paths of the Baso-E cell type after *GATA1* knockout network, showing the 20 perturbation-only genes with the highest path score. **d**, Overlap of perturbation-only peaks with ChIP binding of the perturbed factor in the ten top-ranked of all contexts, compared with matched random peak sets. **e**, A perturbation-only link from an intergenic peak to *SEMA4B* after *GFI1B* knockout in Baso-E, with RNA, accessibility, ChIP and GATA1 HiChIP tracks in control (grey) and knockout (red) cells.

We first compared NT and GATA1-knockout cells at the Baso-E stage. A stratified rank-permutation test of gene attribution classified 403 genes as perturbation-favored and 301 as control-favored, whereas 5,234 genes showed no significant difference (**Fig. 5b**). Perturbation-favored genes were enriched for genes that are down-regulated after GATA1 knockout in observed cells (110/403; 1.94-fold; *p* = 4 × 10⁻¹³). In contrast, control-favored genes were depleted of them (8/301; 0.19-fold; *p* = 6 × 10⁻¹²). For the 20 perturbation-only genes with the highest path score (attention × gene attribution × peak attribution), all 32 routes and 60 element-peak edges were also present in the control network, and only their 23 element-gene edges were perturbation-only (**Fig. 5c**). All 218 perturbation-only genes were reachable through such peak-element-gene paths (**Supplementary Fig. 17**).

Perturbation-only peaks were enriched at binding sites of the perturbed factor. After GATA1 knockout, 59% of the knockout-only peaks overlapped GATA1 ChIP-seq signals (*p* = 1 × 10⁻⁵; **Supplementary Fig. 18**). More broadly, peaks attributed only in the perturbation network overlapped with ChIP-defined binding sites of the perturbed factor significantly in 43 different training contexts. The top-10 ranked contexts were illustrated in **Fig. 5d** and eight of the ten top-ranked contexts involved knockouts of *GATA1*, *TAL1*, *LDB1* or *LMO2*, which act together in a core erythroid transcriptional complex ^39^.

As an example of a perturbation-only link supported by binding and contact data, after *GFI1B* knockout an intergenic peak bound by GATA1 and TAL1 entered the attributed peaks of ATAC element A125 and was thereby linked to *SEMA4B* through the A125-R166 route, which is present in both states. In GATA1 HiChIP, the peak bin contacted the *SEMA4B*-201 promoter, and the peak lost accessibility in real knockout cells, as the model predicted (**Fig. 5e**; the selection rule is given in **Supplementary Fig. 19**). A peak at the *SEMA4B* promoter, also bound by GATA1 and TAL1, reached *SEMA4B* through a second route, A156-R166, in both states, at a cut-off of 500 peaks per element. Together, these observations identify a GFI1B-sensitive distal region connected to *SEMA4B* within GATA1/TAL1-bound chromatin. They support a model in which *GFI1B* helps maintain accessibility at this distal regulatory element, while the promoter-associated route remains represented in both model states. The perturbation-only link therefore pinpoints a regulatory region affected by *GFI1B* loss, rather than simply marking a newly accessible site. Another perturbation-only association involved a peak 1.4–2.3 kb upstream of the *MAGI3* TSS that overlapped reference ChIP-seq peaks for GFI1B, GATA1 and TAL1. This peak was linked to *MAGI3* in the *GFI1B*-knockout network but not in the control network. Both peak accessibility and *MAGI3* expression decreased in observed knockout cells, consistent with the predicted directions of change (**Supplementary Fig. 20**).

Together, these results indicate that TF knockouts preserve the attention backbone but re-weight its features toward genes that respond to the knockout and toward regulatory elements bound by the perturbed factor or its partners, and that single perturbation-only links can be traced to factor-bound elements, such as the enhancer that contacts the *SEMA4B* promoter. Beyond predicting how perturbations reshape paired multiomic states, MultiFlow therefore provides an interpretable map of candidate regulatory elements and target genes through which these responses may propagate, yielding hypotheses that can be tested experimentally.

## Discussion

MultiFlow is built on the premise that a cellular response to perturbation is a coordinated transition across molecular layers. By coupling RNA and ATAC flows within a unified generative process, MultiFlow first learns how gene expression and chromatin accessibility states are jointly organized and then models how perturbation reshapes this paired state across cellular contexts. This formulation connects multiomic generation with perturbation prediction and provides a natural extension of single-cell perturbation modeling beyond transcriptional responses alone. Across our analyses, MultiFlow preserved cell-type-specific multiomic distributions, generalized perturbation responses to unseen cellular contexts, and maintained coordination between predicted transcriptional and chromatin-accessibility changes. Together, these results support a broader view of perturbation prediction, which involves the coordinated change across multiple regulatory layers.

Several limitations exist in the present study. First, although multiomic generation was evaluated across independent datasets, multiomic perturbation prediction was primarily validated using the Perturb-multiome resource ^14^. The extent to which MultiFlow generalizes across experimental platforms therefore remains to be established. This limitation also reflects the current experimental landscape where large paired multiomic perturbation datasets remain relatively scarce ^13,40^.

The second limitation is that the current MultiFlow architecture is designed specifically for two paired modalities. Emerging perturbation assays are likely to measure increasingly rich combinations of transcriptomic, epigenomic, proteomic and other molecular states within the same cells ^5,14,41,42^. Extending MultiFlow to such settings will require a more general multi-flow architecture that can coordinate multiple modality-specific dynamics while remaining computationally scalable and robust to partially observed modalities.

The third limitation concerns the interpretation of the learned cross-modal relationships. The current cross flow modules (CFMs) are data driven, and the RNA-ATAC relationships recovered by MultiFlow are therefore primarily associative. Concordance between accessibility changes and nearby transcriptional responses, or preservation of RNA-ATAC neighborhood structure, does not establish a causal regulatory mechanism. Bridging this gap represents an important direction for future, potentially through the incorporation of causal structural learning ^43,44^ directly into the cross-modal mechanism.

Looking forward, MultiFlow provides a foundation for increasingly general models of multiomic cellular responses. As perturbation datasets expand in scale, molecular coverage and biological diversity, future versions of the framework could integrate additional modalities, richer perturbation representations and causal regulatory structure to model how cellular states transition in different cellular contexts under different interventions. Such developments may move single-cell perturbation modeling toward a more comprehensive and mechanistically informed prediction of cellular responses across contexts.

## Methods

### Problem formulation and notation

MultiFlow operates on paired single-cell multi-omics profiles, in which RNA and ATAC measurements are obtained from the same cell. For each cell, let *x^R^*represent the scRNA-seq expression profile, and let *xx^AA^*denote the scATAC-seq chromatin accessibility profile. The objective of MultiFlow is to model the conditional joint distribution of paired RNA and ATAC profiles:

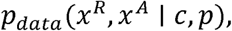

where *c* denotes the cellular context (e.g., cell type), and *p* denotes the perturbation condition. We use *p* = ∅, to denote unperturbed or null condition, whereas *p* ∈ *P* denotes a specific perturbation, such as perturbation of a transcription factor.

To reduce dimensionality, modality-specific autoencoders (*E_R_*, *D_R_*) and (*E_A_*, *D_A_*) are used to project the two modalities RNA and ATAC from original spaces into latent spaces, respectively:

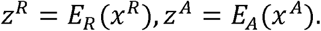

MultiFlow learns the conditional joint distribution in the paired latent space parametrized by *θ*:

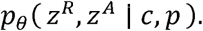

Then generated latent states are mapped back to the original feature spaces through modality-specific decoders,

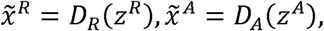

where 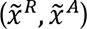 is the multiomic profile generated by MultiFlow.

MultiFlow is designed to support multiomic cell state generation and perturbation prediction within a unified framework. Cell-state generation corresponds to the special case in which the perturbation condition is inactive where *p* = ∅,. MultiFlow generates paired RNA and ATAC states conditioned on cell type *c* under the null condition:

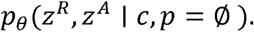

For perturbation prediction, *p* specifies an active perturbation, such as a transcription-factor perturbation. MultiFlow then generates perturbed multiomic cell states conditioned on the cellular context and perturbation condition:

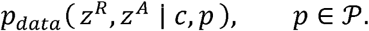

Modeling multiomic cell state generation and perturbation response prediction within a unified framework highlights the flexibility of MultiFlow in multiomic single-cell analysis.

### Latent representation modeling of single-cell multiomics data

MultiFlow models paired RNA and ATAC profiles in modality-specific low-dimensional latent spaces *z* = (*z^R^*, *z^A^*), rather than directly in the original high-dimensional feature spaces. This representation reduces dimensionality and sparsity of single-cell measurements while retaining modality-specific structure that can subsequently be coupled by the flow-matching model. Let *E_R_*, denote the RNA and ATAC encoders, respectively, which map the observed profiles to 128-dimensional latent representations *z^R^*, *z^A^*. Corresponding decoders *D_R_*, *D_A_* map generated latent states back to the original RNA and ATAC feature spaces. Both encoders and decoders adopt multi-layer perceptron (MLP) with three hidden layers. The two modality-specific autoencoders are trained independently before flow-matching training using reconstruction losses in their respective feature spaces ^45–47^. After training, the encoder and decoder parameters are fixed, and MultiFlow operates on the resulting paired latent representations.

### Flow matching generative model

Flow matching is a generative modeling framework ^18^ that learns a continuous transformation between a simple reference distribution (e.g., a standard Gaussian distribution) and the data distribution through a time-dependent vector field. Let *z*_1_∼*p_data_*(*z*) denote a latent representation of an observed cell state and *z*_0_∼*N*(0, *I*) denotes a sample from the Gaussian reference distribution. We define a linear probability path between the two endpoints,

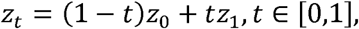

with the corresponding target velocity *u_d_*(*z_d_*|*z*_0_, *z*_1_) = *z*_1_ − *z*_0_. Flow matching learns a neural vector field *v_θ_*(*z_d_*, *t*) to approximate this velocity by minimizing the flow-matching objective

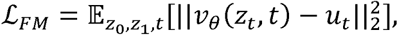

where flow time *t* is sampled uniformly from [0,1]. After training, latent states are generated by sampling random noise from the Gaussian reference distribution and solving the ordinary differential equation

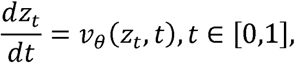

which transports samples from the Gaussian reference distribution at *t* = 0 toward the learned data distribution at *t* = 1. In practice, the trajectory is obtained by numerically integrating the learned vector field from *t* = 0 to *t* = 1 through an ODE solver ^48^. MultiFlow extends this formulation to paired RNA and ATAC latent states by learning two interacting conditional velocity fields, as described below.

### Conditional coupled flow matching

MultiFlow extends standard flow matching to paired RNA and ATAC latent states by learning two modality-specific but interacting data flows. Let 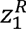 and 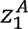 denote the latent representations of an observed paired multiomic cell. The two data flows start from a shared Gaussian sample *z*_0_ ∼ *N*(0, *I*) and are interpolated toward their respective target states,

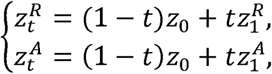

where *t* ∈ [0,1]. The corresponding target velocities are

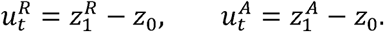

Rather than learning the two flows independently, MultiFlow jointly learns their velocity fields so that the update of each modality depends on the evolving states of both modalities. Specifically,

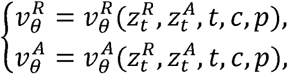

where 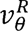 and 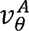 are the RNA and ATAC velocity fields learned by MultiFlow, respectively. *c* represents the cellular context and *p* denotes the perturbation condition. Cross-modal information is exchanged between the two velocity fields through the Cross Flow Modules described below. This design preserves modality-specific trajectories while allowing the RNA and ATAC states to evolve as a coupled generative system.

The two velocity fields are jointly trained using

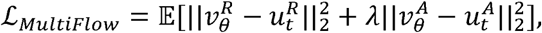

where *λ* controls the relative contribution of the ATAC flow-matching objective and is set to 1 in all experiments.

At inference, paired latent states are generated by jointly integrating the learned RNA and ATAC velocity fields from *t* = 0 to *t* = 1 by solving the two coupled ODEs along their learned trajectories. These generated latent vectors are then decoded back to the original RNA and ATAC feature spaces using the corresponding AE/VAE decoders.

#### Cellular context representation

MultiFlow uses task-specific representations of cellular context for cell-state generation and perturbation prediction tasks. For cell-state generation, the context for cell type *c* is represented by a learnable embedding of the cell-type label. This representation provides the categorical cellular identity used to condition generation under the null perturbation.

For perturbation prediction, cellular context embedding is instead derived directly from the unperturbed multiomic state of the cell population, allowing the model to represent cellular contexts whose perturbed states are not observed during training. Specifically, let ℐ*_c_* = {*i*: *p_i_* = ∅,} denote the cell index set for unperturbed cells of cellular context *c*. Then, the control-derived context representation is defined as the mean of the paired RNA and ATAC latent states,

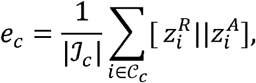

where || denotes concatenation. This representation summarizes the baseline transcriptional and chromatin accessibility state of the cellular context *c* and can be constructed directly from observed control cells.

To evaluate generalization ability of MultiFlow to predict perturbation responses in unseen cellular contexts, we used a leave-one-cell-type-out setting. For each held-out cell type, the perturbed cells are excluded from training, thereby avoiding information leakage from the held-out perturbed responses during training. Control cells from the held-out cell type are used only to define the baseline cellular context embedding during inference.

#### Perturbation representation

We denote the perturbation condition by *p*, where *p* = ∅, represents the unperturbed control condition and *p* ∈ *P* denotes a specific active perturbation. Here, *P* ={*p*(1), …, *p*(K)} represent *K* distinct perturbation conditions. In the perturb-multiome experiments, each active condition corresponds to perturbation of an individual transcription factor.

For perturbation-response prediction, each active perturbation *p* is mapped to a learnable embedding, denoted as= *E_emb_*(*p*). The embedding mapping is learned jointly with the flow matching model.

For cell state generation, the null condition *p p* = ∅, represents the absence of an active perturbation, the perturbation conditioning channel is therefore kept inactive and represented by an all-zero vector. This allows a flexible modeling in MultiFlow to tackle two different tasks without modifying the model architecture.

### MultiFlow architecture

MultiFlow parameterizes the coupled RNA and ATAC velocity fields using a dual-branch residual network with bidirectional cross-modal information exchange. At flow time *t*, the RNA and ATAC latent states 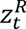 and 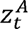, are independently projected from their 128-dimensional latent spaces into a common *d*_ℎ_ = 512 dimensional hidden space, denoted as 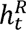 and 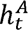, respectively. The two hidden representations are subsequently processed by parallel residual branches with U-Net ^49^ style skip connections. Cross-modal information is exchanged at multiple depths through bidirectional Cross Flow Modules (CFMs), while RNA and ATAC remain represented by separate hidden states throughout the network. The flow time t is represented as *e_d_* using a sinusoidal embedding, followed by a two-layer multilayer perceptron, where both hidden and output dimensions are 512. Finally, the time embedding, context embedding, and perturbation embedding are normalized separately and then added together to form a combined conditional embedding vector,

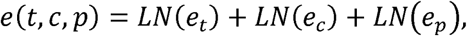

where LN denotes LayerNorm ^50^. Context embedding *e_c_* and perturbation embeddin are defined in previous section. We duplicated e*_p_*∈ ℝ^256^along the feature dimension to match the 512-dimensional network representation before normalization. The context representation is fixed during flow-model training, whereas embeddings of active perturbations are learned jointly with the velocity fields.

The resulting conditional vector is incorporated throughout the network through condition-modulated residual blocks. Let ℎ ∈ ℝ^512^denote the RNA or ATAC hidden representation entering a residual block. The hidden state is first normalized and transformed through a SiLU-activated linear layer to obtain an intermediate representation *u*. In parallel, the condition vector is projected to feature-wise scale and shift parameters ^49,51^, which modulate the normalized intermediate representation as,

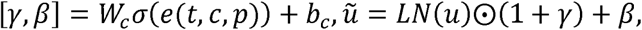

where *σ* denotes the SiLU activation and *ẋ* denotes the element-wise product. The modulated representation ũ is subsequently passed through a second SiLU-activated linear transformation and added to the input h through a residual connection. In this way, flow time, cellular context and perturbation modulate the evolving RNA and ATAC representations at each residual stage, rather than being introduced only at the network input.

Overall, the network consists of two encoder residual blocks, one middle residual block, two decoder residual blocks and three bidirectional Cross Flow Modules (CFMs), which are introduced below.

#### Cross Flow Module

The Cross Flow Module (CFM) enables cross modality information exchange through a bidirectional cross-attention module between RNA and ATAC hidden features. For a given cell, the RNA and ATAC hidden states before entering a CFM are denoted as ℎ*^R^*, ℎ*^A^* ∈ ℝ*^d^*^ℎ^, respectively (*d*_ℎ_ = 512). MultiFlow treats the *d*_ℎ_ hidden coordinates as feature tokens. Each scalar hidden feature is projected into an attention space of dimension *d_a_*= 64, producing *Q_R_*, *K_R_*, *V_R_* ∈ ℝ*^d^*^ℎ×*d*^*^a^*, and *Q_A_*, *K_A_*, *V_A_* ∈ ℝ*^d^*^ℎ×*d*^*^a^* in the attention mechanism ^19^.

Attention from RNA features to ATAC features is computed as a cross-attention matrix:

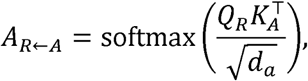

and the RNA representation is updated as 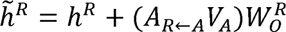. Conversely, we have

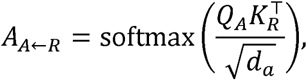

and the ATAC representation is updated as 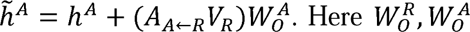 project the 64-dimensional attention outputs back to scalar hidden features. The attention matrices above have the size 512 × 512, allowing individual hidden features in one modality to selectively integrate information from the other modality. Biologically, the cross-attention allows the model to represent coordinated changes in transcription and chromatin accessibility while retaining modality-specific response patterns. All query, key, value and output projections are modality specific.

The complete backbone contains two residual blocks in each branch before the central interaction stage. Their outputs are retained as modality-specific skip representations. The RNA and ATAC states then pass through two consecutive CFMs followed by a middle residual block. In the second half of the network, the deepest skip representation is added back to each modality, followed by a third CFM and a residual block; the remaining skip representation is then incorporated before the final residual block.

Finally, separate linear output heads project the 512-dimensional hidden representations to modality-specific velocity vectors 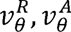 in their respective latent spaces. These vectors specify the instantaneous RNA and ATAC velocities used to evolve the paired latent states along the coupled flow trajectories.

### Perturbation-specific debiasing across cellular contexts

Generative perturbation models may retain systematic prediction bias when transferring responses to an unseen cellular context. We therefore introduce a perturbation-specific debiasing procedure that estimates residual prediction bias from training cellular contexts and transfers this correction to the held-out context. This strategy is inspired by the debiasing principle introduced in dbDiffusion ^52^, which uses prediction errors from observed perturbations to calibrate predictions for an unseen perturbation. Our setting differs in the axis of generalization: the perturbation itself is observed during training, whereas its response in the target cellular context is unseen. We therefore estimate bias across observed cellular contexts for the same perturbation, rather than across related perturbations.

Specifically, for each perturbation *p*, let *c_train_* denote cellular context set during training. For each training context *c*, we compute the mean residual between the observed and MultiFlow-predicted perturbed latent states as:

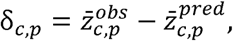

where 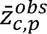 denotes the mean latent representation of observed true cells of cellular context *c* under perturbation *p*, and 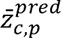 denotes the mean latent representation of generated cells of the same cellular context and under the same perturbation condition.

The perturbation-specific bias transferred across cellular contexts is then estimated as

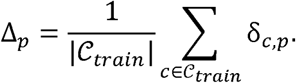

Δ*_p_* represents the perturbation-specific prediction bias of MultiFlow in the latent space. Note that in the leave-one-cell-type-out setting, Δ*_p_* is estimated only from the training data to avoid information leakage.

For a held-out cellular context *c*^′^, the predicted latent state under perturbation *p* is corrected by adding this perturbation-specific correction vector:

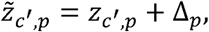

where *z_c_*′_,*p*_ and 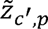 denote the original and debiased MultiFlow predictions, respectively. Importantly, Δ*_p_* is estimated exclusively from the training cellular contexts, and perturbed cells from held-out cellular contexts are never used for bias estimation to avoid information leakage.

### Datasets

We used three paired single-cell multi-omics datasets to evaluate MultiFlow, with the OpenProblems ^23^ and PBMC10k datasets ^25^ used for cell-state generation and the Perturb-multiome dataset (GEO: GSE274113) ^14^ used for perturbation response prediction.

#### OpenProblems multiome dataset

The dataset was used as the primary benchmark for the paired cell state generation task. This dataset contained paired scRNA-seq and scATAC-seq profiles from the same cells, which simultaneously measured the transcriptional states and chromatin accessibility across diverse immune and hematopoietic populations. After preprocessing, the dataset comprised 69,249 cells from 22 annotated cell types, with 13,431 RNA genes and 36,553 ATAC peaks. This dataset was used to evaluate the ability of MultiFlow to recover cell-type-specific RNA and ATAC joint states and their population distributions.

#### PBMC10k multiome dataset

The PBMC10k dataset was used as an independent validation dataset for cell-state generation. After preprocessing, the dataset contained 12,012 cells, from 13 immune-cell populations, including CD4+ T cells, CD8+ T cells and B cells, represented by 13,431 RNA genes and 36,553 ATAC peaks. PBMC10k was used to assess whether the generative performance observed on OpenProblems generalized to an independent paired multiomic dataset.

#### Perturb-multiome dataset

The Perturb-multiome dataset was used for evaluating perturbation response prediction across cellular contexts. It contained paired single-cell RNA and ATAC profiles with perturbation labels across hematopoietic differentiation. After preprocessing, the dataset contained 133,022 cells, spanning 10 annotated hematopoietic cell types, 19 transcription-factor perturbations, and an unperturbed control condition. This dataset was used in the leave-one-cell-type-out evaluation of perturbation responses in cellular contexts whose perturbed states were not observed during training.

### Data preprocessing

RNA and ATAC modalities were preprocessed separately while preserving cell-level pairing between the two modalities. Cells with low read coverage and genes detected in too few cells were removed. Cell type annotations and perturbation labels were retained and used as conditional labels for downstream modeling.

#### OpenProblems multiome dataset

For scRNA-seq profiles, cells with fewer than 100 detected genes were removed, and genes detected in fewer than three cells were filtered using Scanpy ^53^. RNA counts were library-size normalized and log-transformed, after which 13,431 genes were retained for downstream modeling. For scATAC-seq profiles, only cells with matched RNA profiles were retained, and then kept the top 36,553 peaks ranked by the number of cells in which each peak was detected. The scATAC-seq matrix was binarized, with values set to 1 when the read count is greater than or equal to 1 and 0 otherwise. For the generation benchmark, cells were randomly partitioned into 80% training and 20% held-out test cells using a fixed random seed. The same split was used across MultiFlow and all competing methods.

#### PBMC10k multiome dataset

The PBMC10k multiome dataset was preprocessed using the same workflow as the OpenProblems multiome dataset. RNA profiles were filtered, log-normalized, and represented using the selected 13,431-gene feature space. ATAC profiles are restricted to cells with matched RNA profiles, represented using the selected 36,553-peak feature space and binarized. Cells were divided into 80% training and 20% held-out test sets using a fixed random seed, with identical partitions used for all evaluated methods.

#### Perturb-multiome dataset

For the scRNA-seq modality, raw reads were processed using the standard pipeline provided in the original study, and the 14 experimental replicates were then combined and further processed using a Scanpy ^53^ workflow, including cell filtering, log-normalization, and selection of highly variable genes, resulting in 13,431 genes. For the scATAC-seq modality, peaks identified across replicates were merged across replicates based on both reciprocal overlap (>85%) and minimum replicate was represented in more than 10 of the 14 replicates. The resulting peak set was ranked by detection frequency, and the top 36,553 peaks were finally retained for downstream analysis.

For perturbation-response prediction, we used a leave-one-cell-type-out setting. In each fold, one of the ten cell types was designated as the target context. All cells from that cell type carrying an active perturbation were excluded from model training, whereas the remaining nine cell types provided the observed perturbation responses used for training. Unperturbed control cells from the held-out cell type were used to define the target cellular-context representation only in the prediction stage.

### MultiFlow training details

MultiFlow was trained separately for cell-state generation and perturbation-response prediction using the conditional flow-matching objective described above. All modality-specific autoencoders were pretrained before flow-model optimization and kept fixed during flow training.

For the cell-state generation task, the MultiFlow model was trained for 600 epochs with a batch size of 512 using the Adam optimizer ^54^ with a learning rate of 1 × 10^−4^. During each training iteration, paired RNA and ATAC latent states were sampled together with random Gaussian noise and random flow times *t* ∼ *U*(0,1), and the coupled velocity fields were optimized to match the target transport directions. During inference, paired RNA and ATAC states were generated by integrating the learned coupled ordinary differential equations (ODEs) from *t* = 0 to *t* = 1 using a midpoint solver with 100 integration steps. Generated latent states were subsequently mapped back to the original RNA and ATAC feature spaces using the pretrained decoders.

For the perturbation response prediction task, the model was trained for 1600 epochs with the same optimizer, learning rate and batch size. The perturbation-conditioned velocity fields were learned using control-derived cellular-context representations and perturbation embeddings defined above. During inference, perturbed latent states were generated by solving the coupled ODE system using 50 midpoint integration steps. The generated latent representations were then corrected using the perturbation-specific debiasing procedure and decoded into RNA and ATAC profiles for downstream evaluation.

MultiFlow was implemented in PyTorch ^55^ and was run on a single NVIDIA B200 GPU for training and inference.

### Benchmark Methods

For cell-state generation, MultiFlow was compared with CFGen (https://github.com/theislab/CFGen)^21^, MultiVI (https://github.com/scverse/scvi-tools)^22^, scDiffusion-X (https://github.com/EperLuo/scDiffusion-X)^20^, and scDesign3 (https://github.com/SONGDONGYUAN1994/scDesign3)^24^. All methods used the same training and held-out test splits and the same cell-type labels. RNA outputs were evaluated after library-size normalization to 10,000 counts per cell followed by log transformation, whereas ATAC outputs were evaluated as binary accessibility profiles.

For perturbation prediction, MultiFlow was compared with scGen (https://github.com/theislab/scgen)^56^, trVAE (https://github.com/theislab/trvae)^7^, scPreGAN (https://github.com/JaneJiayiDong/scPreGAN)^9^, CellOT (https://github.com/bunnech/cellot)^10^, BioLORD (https://github.com/nitzanlab/biolord)^12^, inVAE (https://github.com/theislab/inVAE)^8^, TxPert (https://github.com/valence-labs/TxPert)^57^, and MultiPert (https://github.com/MengyuanZhaoo/MultiPert)^58^. Implementations available through scPerturBench were adapted from its benchmark repository (https://github.com/bm2-lab/scPerturBench)^6^. TxPert and multipert were implemented from its official repositories. Every method used the same leave-one-cell-type-out split: nine cell types were used for training and the remaining cell type was held out for testing.

Three additional baselines from scPerturBench were implemented for both RNA and ATAC. For baseReg, a separate ridge-regularized scalar regression was fitted for every feature and perturbation across the training cell types. Suppose *x_c_*_,*p*,*j*_ is the perturbed pseudo-bulk value and *x_c_*_,0,*j*_ is the control value, the model was fitted with ridge penalty *λ* = 10^−4^:

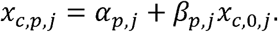

For baseMLP, the control pseudo-bulk profile was concatenated with a one-hot perturbation vector and passed through one fully connected hidden layer with 256 rectified linear units. The output layer predicted all RNA genes or ATAC peaks. The model was optimized by Adam for 200 full-batch epochs using mean squared error and a learning rate of 1 × 10^−3^. TrainMean was calculated by the cell-count-weighted mean perturbed profile across the training cell types.

### Evaluation metrics

Metrics were calculated for both RNA and ATAC modalities. RNA was evaluated in the log-transformed space, whereas ATAC was evaluated using pseudo-bulk accessibility probabilities derived from binary profiles. Let *x_i_*_,*j*_*^e^* and 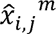 denote the real and generated or predicted values for cell *i* and feature *j*, where *u* could be either *RNR* or *ATAC*. The real and generated sets were sampled with equal cell numbers before calculating the metrics.

Two major categories of metrics were used: mean-profile metrics, which compare feature-wise mean values across bulk cells, and distribution-level metrics, which compare the distributions of individual cells.

#### Mean-profile metrics

For each feature, the real and generated mean profiles were

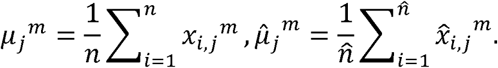

Pearson correlation coefficient (PCC) was calculated between the two feature-mean profiles:

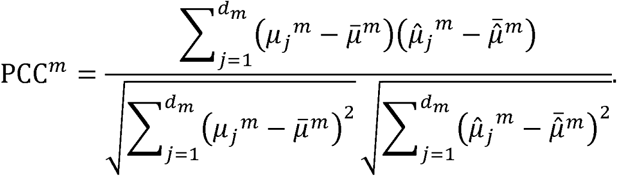

Here, *d_m_* is the number of features in modality *u*.

Spearman correlation coefficient (SCC) was calculated by replacing each profile with its feature ranks:

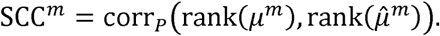

Mean squared error (MSE) was calculated by the mean squared difference between feature means:

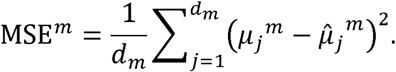

#### Distribution-level metrics

Maximum mean discrepancy (MMD) ^29^ compared the real and generated cell distributions after projecting them into a common principal-component analysis (PCA) space ^59^. For real embeddings *U* = {*u_i_*} and generated embeddings 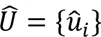, MMD is,

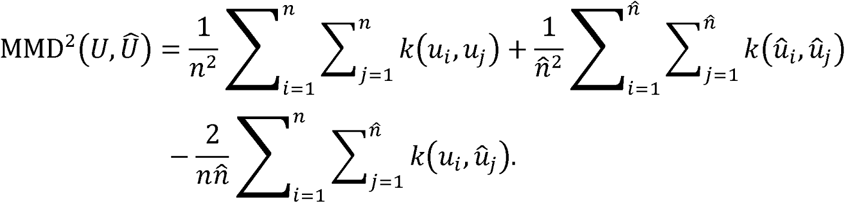

For cell-state generation, we followed the scDiffusion-X implementation: the kernel was the sum of five Gaussian kernels with bandwidth multiplier 2, and the diagonal terms were retained. For pooled MMD, cells were combined across all cell types, and equal numbers of real and generated cells were sampled. For perturbation prediction, MMD was calculated on the top 100 response features with the largest perturbation effects, and was therefore referred to as top-100 MMD.

Local inverse Simpson’s index (LISI) ^27^ measured local mixing between real and generated cells in the same common PCA space. For cell *i*, let *p_i_*_,*e*_ be the fraction of its nearest neighbors assigned to batch *b*, where the two batches were real and generated. The local score was

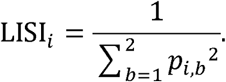

We used the scIB-scaled integration LISI (iLISI) score on the interval from 0 to 1, where larger values indicate stronger real-generated mixing. The generation benchmark used 10 nearest neighbors and the first 20 principal components of PCA, following the scDiffusion-X setting. The perturbation benchmark used the same setting as generation benchmark.

#### Perturbation-effect metrics

For cell type *c* and perturbation *p*, the real and predicted perturbation effects were calculated by:

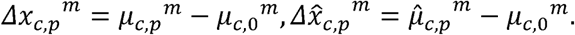

where *μ_c,p_^m^* represents the mean profile of cells of type *c* under perturbation *p*, and *μ_c_*_,0_*^e^* represents the mean profile of the corresponding control cells.

The top 100 perturbation-responsive genes were ranked within each held-out context from the real perturbation effect. PCC-delta was calculated as,

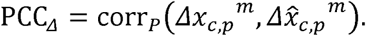

Delta MSE was

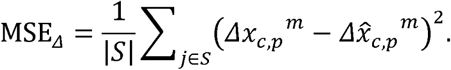

Here, *S* is the selected feature set, where we used the top-100 genes with the largest perturbation effects. Scores were first computed for each held-out cell-type-perturbation and were then averaged across cell-types.

#### ATAC pseudo-bulk metrics

Because single-cell ATAC accessibility profiles were binary, some ATAC metrics were calculated using pseudo-bulk accessibility values. The pseudo-bulk accessibility of peak *j* in cell type *c* and condition *p* was the fraction of cells in which the peak was accessible:

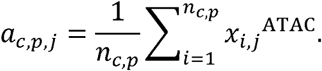

where *n_c,p_* is the number of cells belonging to cell type *c* and perturbation *p*. ATAC perturbation effects were defined as the difference between perturbed and control accessibility,

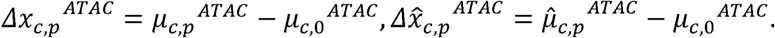

For the ATAC modality, we calculated effect cosine similarity and direction accuracy on a selected peak set *S*, which is,

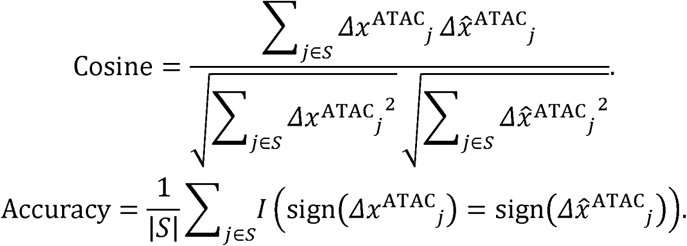

Effect cosine similarity measured the correlation between the real and predicted accessibility-change vectors, whereas direction accuracy measured the fraction of peaks for which the predicted and real accessibility changes in the same direction.

For the significant-peak analysis, the 5,000 most variable peaks were selected within each training fold. Peaks were tested between control and perturbed cells using Welch’s t-test, followed by BH correction. Up to 100 peaks with false-discovery rate below 0.05 were retained per context. Significant-delta Pearson correlation, direction accuracy, iLISI and MMD were then calculated on this peak set. The composite absolute score was the unweighted mean of Pearson correlation, direction accuracy, iLISI and 1-MMD:

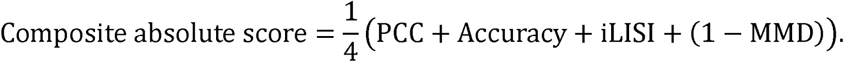

For visualization, real and generated or predicted cells were embedded together after principal-component analysis using UMAP.

### Cell-type identity classification

The cell-type identity matrices were obtained with classifiers trained only on real training cells. For each cell type, at most 500 training cells were sampled. Features were ranked by the average feature variance in held-out real and generated cells, and the top 2,500 RNA genes or 4,000 ATAC peaks were retained. RNA values were evaluated in the normalized log space and ATAC values in the binary space. Truncated singular-value decomposition with 40 components and a standard scaler were fitted to the real training set and then applied unchanged to held-out real and generated cells. A class-balanced multinomial logistic-regression classifier was trained with a regularization parameter C = 2 and a maximum of 2,500 iterations. After classification, the rows of each confusion matrix were normalized to sum to one. The diagonal therefore represents the fraction of cells generated under cell type *c* that were classified as the correct cell type.

### Peak-gene effect concordance

To evaluate whether predicted ATAC effects were consistent with transcriptional effects in nearby genes, we defined the peak-gene effect concordance score as follows. Based on the expectation that peaks near upregulated genes become more accessible, whereas peaks near downregulated genes become less accessible, we calculated peak-gene effect concordance to quantify the agreement between these coordinated RNA and ATAC effects. Gene neighborhoods were defined as ±100 kb around transcription start sites using the Homo sapiens GRCh38.86 annotation ^60^. For every held-out cell-type-perturbation combination, the top 100 RNA genes ranked by the real t-test effect were separated into upregulated and downregulated sets. Peaks with Welch’s t-test *p*-values below 0.05 were considered significant. Significant peaks with positive and negative accessibility changes were classified as opening and closing peaks, respectively.

Significant opening and closing peaks identified from the real data served as reference sets. The same numbers of predicted opening and closing peaks were then selected from each prediction method. The enrichment score was calculated as the proportion of selected peaks located near genes with matched perturbation effects minus the corresponding proportion among background regions. Let 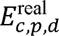 and 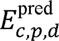 denote the enrichment scores obtained from the real and predicted profiles, respectively, for cell type *c*, perturbation *p*, and effect direction *d*, where *d* could be upregulated/opening or downregulated/closing. Peak-gene effect concordance was defined as the Pearson correlation between the real and predicted enrichment scores:

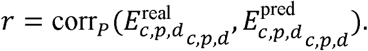

A higher correlation indicates better recovery of coordinated RNA and ATAC perturbation effects.

### Recovery of cell-level RNA-ATAC coordination

To assess whether paired RNA and ATAC predictions preserved cross-modal organization at the single-cell level, 64 paired cells were sampled from each held-out cell-type-perturbation context. RNA and ATAC were reduced separately to 30 principal components. For each cell, 15 nearest neighbors were identified independently in the RNA and ATAC spaces. If 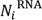 and 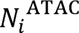 denote these two neighbor sets, their overlap was quantified by the Jaccard index:

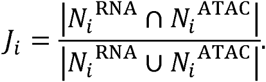

The cell-level Jaccard indices were averaged within each cell type. RNA-ATAC coupling neighborhood-concordance score was the Pearson correlation between the real and predicted context-level mean Jaccard indices. A higher correlation indicates better recovery of coordinated cell-state organization between transcriptional and chromatin-accessibility profiles.

### Attention analysis and feature attribution

All interpretability analyses used the trained MultiFlow models with fixed parameters. We refer to the 512 hidden coordinates of the RNA and ATAC representations that enter a CFM as RNA and ATAC hidden elements. Analyses used the second CFMs (CFM2) at flow time *t* = 0.90 (Supplementary Notes). For a set of *n* cells, we combined the two cross-attention matrices and averaged them across cells:

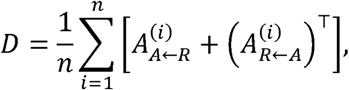

The resulting matrix has ATAC elements as rows and RNA elements as columns. We ranked ATAC-RNA element pairs (*u*, *r*) by their corresponding attention scores *D_d_*_,*t*_.

To associate hidden elements with genes and peaks, we computed gradient-based input attribution. Each hidden element, attribution was defined as the mean absolute gradient with respect to an input feature (a gene or a peak) of the same modality:

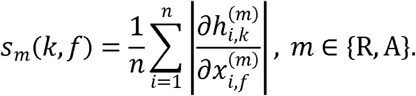

Here, 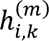 denotes hidden element *k* in modality *u* for cell *i*, and 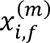 denotes the corresponding input gene-expression or peak-accessibility value. This score measures the sensitivity of a hidden element to an input feature.

For cells generated in the perturbation state, we additionally computed output attribution to quantify the sensitivity of generated gene expression to individual RNA hidden elements:

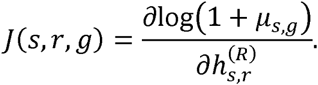

Here, 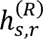 is RNA hidden element *r* in generated sample *s*, and *μ_s,g_* is the decoded expression of gene *g*. Gradients were propagated through the remaining integration steps and the RNA decoder. Combining an attention pair (*u*, *r*) with an attributed peak of ATAC element *u* and an attributed gene of RNA element *r* defined a peak-ATAC element-RNA element-gene path. These paths were interpreted as candidate peak-gene associations within the model. The choice of CFM2 and *t* = 0.90 was guided by comparisons across modules and flow times (**Supplementary Fig. 21** and **Supplementary Note**). In general, attention maps were nearly identical across cell types when flow time *tt* is close to 0, when flow states were dominated by the Gaussian source, and became more cell-type-specific at later times.

### Cell-type markers and regulatory annotations

Within each cell type, genes were scored by averaging input attribution across the 20 RNA elements receiving the strongest attention from ATAC elements. Genes whose summed normalized log-expression across the analyzed cells did not exceed threshold (0.2 times the number of cells) were given a score of zero but kept in the background. Scores for positive marker genes from the original cell-type annotation ^23^ were compared with all other genes by a two-sided Wilcoxon rank-sum test, with BH correction across the 22 cell types.

For genomic overlap analyses, we selected the distinct RNA and ATAC elements represented in the 15 highest-scoring attention pairs. The 100 highest-attribution genes per RNA element and 200 highest-attribution peaks per ATAC element were combined into gene and peak sets. Promoters were defined as the 3-kb regions upstream of gene starts annotated in GENCODE v47 ^61^. We counted attributed genes with at least one promoter-overlapping attributed peak, and overlapping pairs of attributed peaks and lineage-matched H3K27ac intervals^20^. Observed counts were compared with size-matched random gene and peak sets using a one-sided one-sample *t*-test against 20 random sets. Empirical *p*-values were additionally calculated from 1,000 random sets and adjusted using the BH procedure.

### Gene modules and network enrichment

Using the Perturb-multiome model, we computed output attribution for all 512 RNA elements. Absolute attribution values were normalized within each sample and element, then averaged across samples. For each gene, element-specific scores were compared with 99,999 random scores, each random score constructed by selecting one RNA element’s attribution scores at random in each sample. Genes with empirical *p*-value<0.0005 were assigned to the corresponding element, retaining at most the 200 highest-scoring genes for further analysis.

Functional enrichment was assessed using human GO biological-process gene sets from MSigDB C5 ^62^. One-sided hypergeometric tests used the tested genes as background, with BH correction within each perturbation-cell-type combination. Similarity between element-associated gene sets was measured by the Jaccard index, and similarity between their GO enrichment profiles was measured by the cosine similarity of −log_10_ *p*-values across all tested terms.

To construct gene regulatory networks, two genes were linked when assigned to the same RNA element, and each link was weighted by the number of elements that selected the pair. For the 20 training contexts with the highest RNA PCC-delta, we counted the gene pairs supported by TRRUST v2 ^34^ or STRING v12 ^35^ using functional score ≥ 400 or physical score ≥ 700. To obtain the number expected by chance, we kept the network unchanged and only shuffled the gene names.

Enrichment was assessed against 999 permutations of gene labels, in which each gene was swapped only with genes selected by the same number of elements, so that the enrichment cannot be explained by frequently selected genes alone. This preserves network topology and gene-selection frequencies. Enrichment *p*-values were adjusted using the BH procedure.

To find candidate link not yet in reference databases, we prioritized gene pairs selected together by at least three RNA elements. Because frequently selected genes often appear together by chance, each pair also had to pass a one-sided hypergeometric test (*p* < 0.01) based on how often each of the two genes is selected. These pairs were then examined for reference support. In Fig. 4i, a selected pair was considered known when it had a TRRUST record or a STRING score ≥ 150 (functional or physical). And pairs with no record in either database were reported as new candidate edges.

### Comparison of control and perturbation networks

For each training context of the Perturb-multiome data, we passed the observed non-targeting control cells and the knockout cells through the trained model and built one gene regulatory network for each condition. A network links peaks to ATAC elements, ATAC elements to RNA elements, and RNA elements to genes by gradient-based strategy. Each condition’s network comprised the 1,000 highest-scoring attention pairs and the 100 highest-attribution features per participating element.

Genes, peaks and peak-gene links present only in the knockout network but not in the control network were called perturbation-only. To rank perturbation-only genes, an attention pair (*a*, *r*) linked to gene was scored as the product of three values: the attention between *a* and *r*, the attribution of the gene to RNA element *r*, and the attribution of the top-ranked peak of ATAC element *a*.

Perturbation-only peaks were compared with binding-site annotations for the perturbed factor, using one reference dataset selected in advance for each factor. These references comprised ChIP-seq, ChIPmentation ^63^ and CUT&RUN peak sets ^64^. Overlap enrichment was assessed against 99,999 random peak sets matched for chromosome, accessibility and peak width, with BH correction across perturbation-cell-type combinations. Chromatin-contact support was assessed using GATA1 HiChIP from CD34⁺-derived erythroid cells. The published FitHiChIP loops had 5-kb resolution and were converted from hg19 to hg38. A peak-gene link was considered contact-supported when the peak and a transcription start site of the gene overlapped opposite loop anchors.

For the CD14⁺ monocyte network in multiome dataset, we used the 263 attention pairs in the top 0.1% of CD14⁺ monocytes, with attributed genes, peaks and promoters defined as above. A pair linked a peak to a gene when one of the attributed peaks of its ATAC element overlapped both the promoter of one of the attributed genes of its RNA element and a CD14⁺ monocyte H3K27ac ChIP-seq interval. Each route was scored by the product of the attention of the pair, the attribution of the gene and the attribution of the peak. The 20 genes with the highest scores, each with up to three routes were shown in the figure. GO enrichment of all genes in the network used the 13,431 measured genes as background.

## Data availability

The raw data from the perturbation study are available from the Gene Expression Omnibus under accession GSE274113. The processed multiomic H5MU file used for the perturbation analyses can be found in Zenodo at https://doi.org/10.5281/zenodo.21986866. The OpenProblems paired multiome dataset used for cell-state generation study is available from Figshare at https://doi.org/10.6084/m9.figshare.28582061.v3. The PBMC10k multiome dataset used for cell-state generation study is available from 10x Genomics at https://www.10xgenomics.com/datasets/pbmc-from-a-healthy-donor-no-cell-sorting-10-k-1-standard-2-0-0. Public reference data used in the interpretation analyses are listed with their accession numbers and versions in Supplementary Table 1.

## Code availability

The MultiFlow source code, installation instructions and tutorials are available at https://github.com/liuq-lab/MultiFlow.

## Acknowledgements

H.C.W. and Q.L. were partially supported by the National Institutes of Health (NIH) grant R00HG013661. Q.L. was partially supported by Operations Core of the Claude D. Pepper Older Americans Independence Center at Yale School of Medicine (P30AG021342), and the YSPH Transformation Pilot Grant.

