## Supplementary File for "MultiFlow: coupled flow matching for predicting single-cell multiomic perturbation responses in unseen cellular contexts"

### Supplementary Information

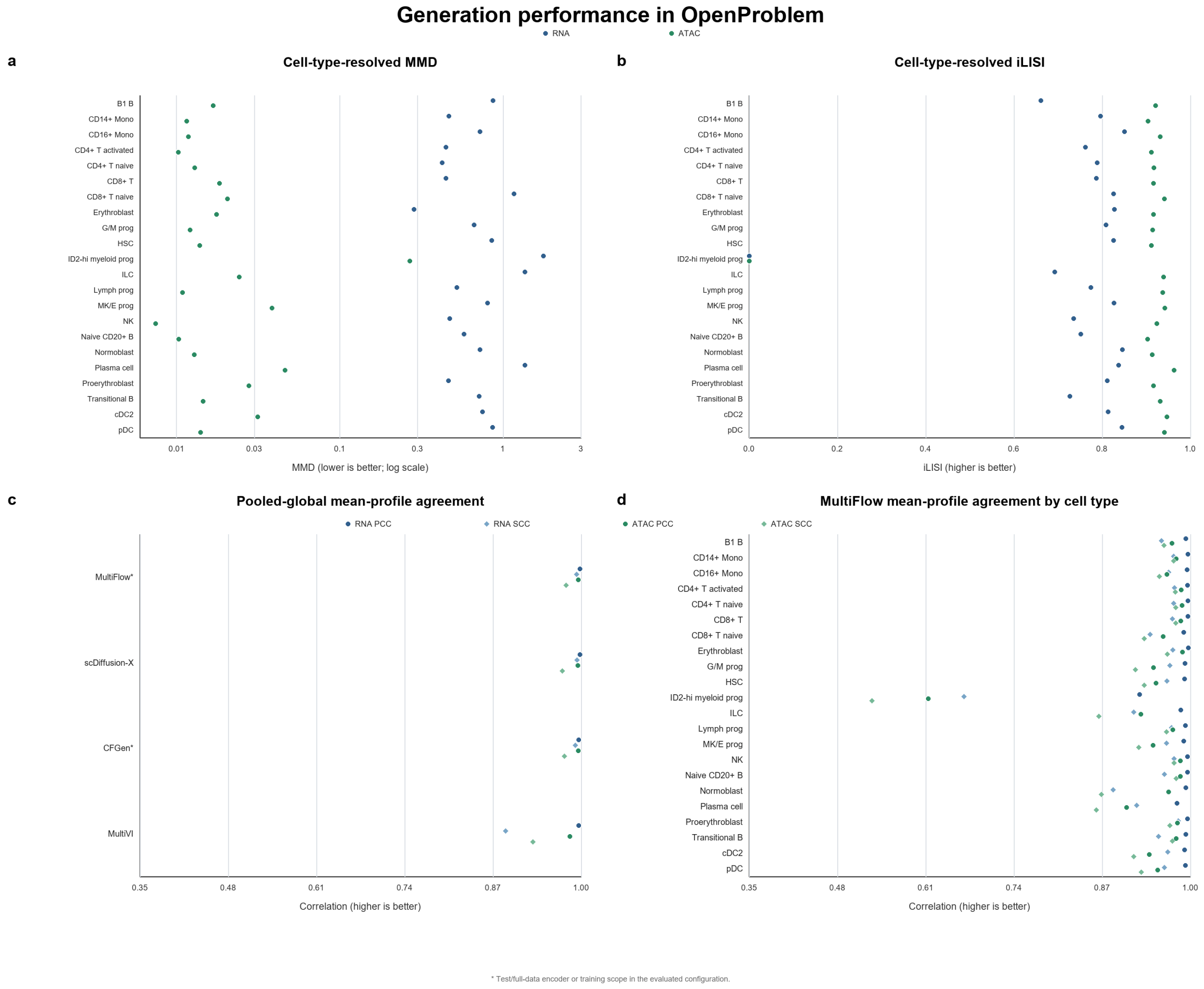


**Supplementary Figure 1.** Generation performance in OpenProblem. a and b, Cell-type-resolved maximum mean discrepancy (MMD) and integration local inverse Simpson's index (iLISI) for MultiFlow-generated RNA and ATAC profiles. Each point denotes one cell type. MMD is shown on a logarithmic scale; lower MMD and higher iLISI indicate better agreement between real and generated distributions. c, Pooled-global Pearson correlation coefficient (PCC) and Spearman correlation coefficient (SCC) across generation methods. d, Cell-type-resolved PCC and SCC for MultiFlow. Circles denote PCC, diamonds denote SCC, and colors distinguish RNA from ATAC; higher values indicate stronger agreement with real mean profiles.

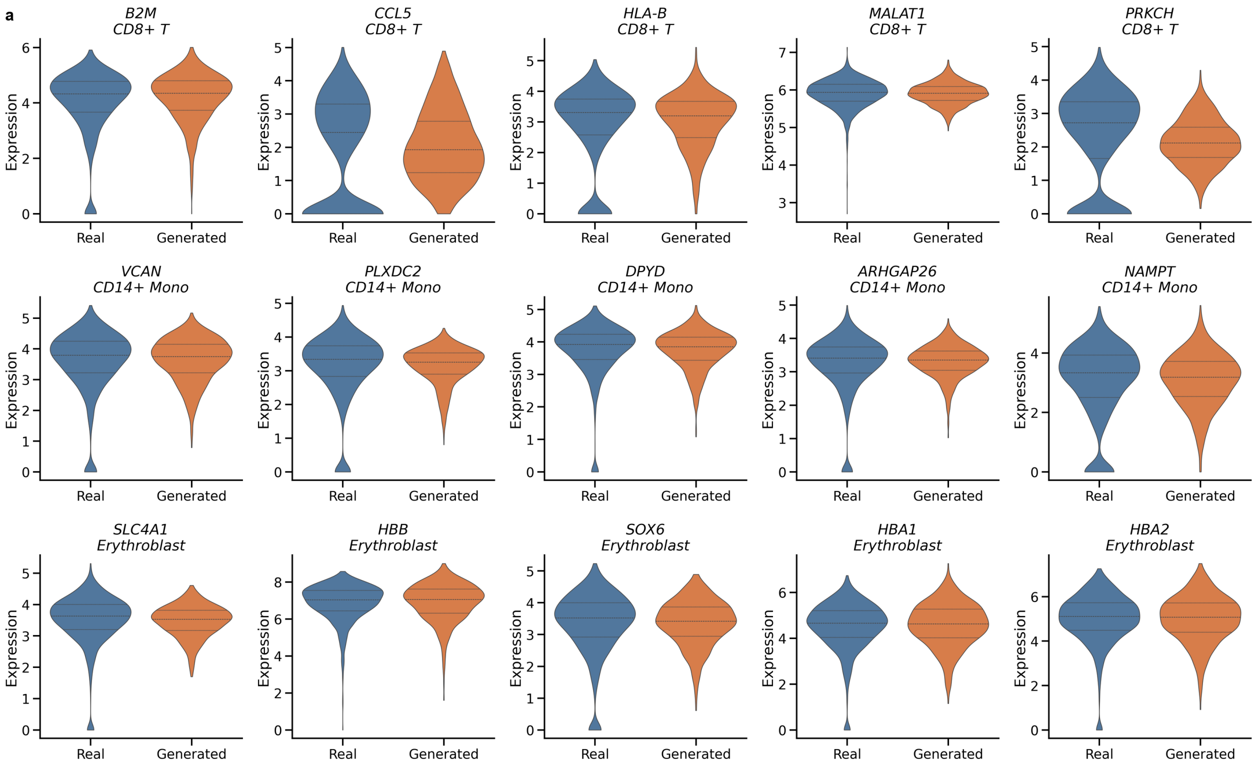


**Supplementary Figure 2.** Cell-type-specific gene-expression distributions in real and generated cells. a, Distributions of five top-ranked RNA differential genes in each of three representative cell types: CD8+ T cells (B2M, CCL5, HLA-B, MALAT1 and PRKCH), CD14+ monocytes (VCAN, PLXDC2, DPYD, ARHGAP26 and NAMPT), and erythroblasts (SLC4A1, HBB, SOX6, HBA1 and HBA2). Violin plots compare held-out real and MultiFlow-generated cells; horizontal lines denote quartiles. Expression is shown on the log-normalized scale.

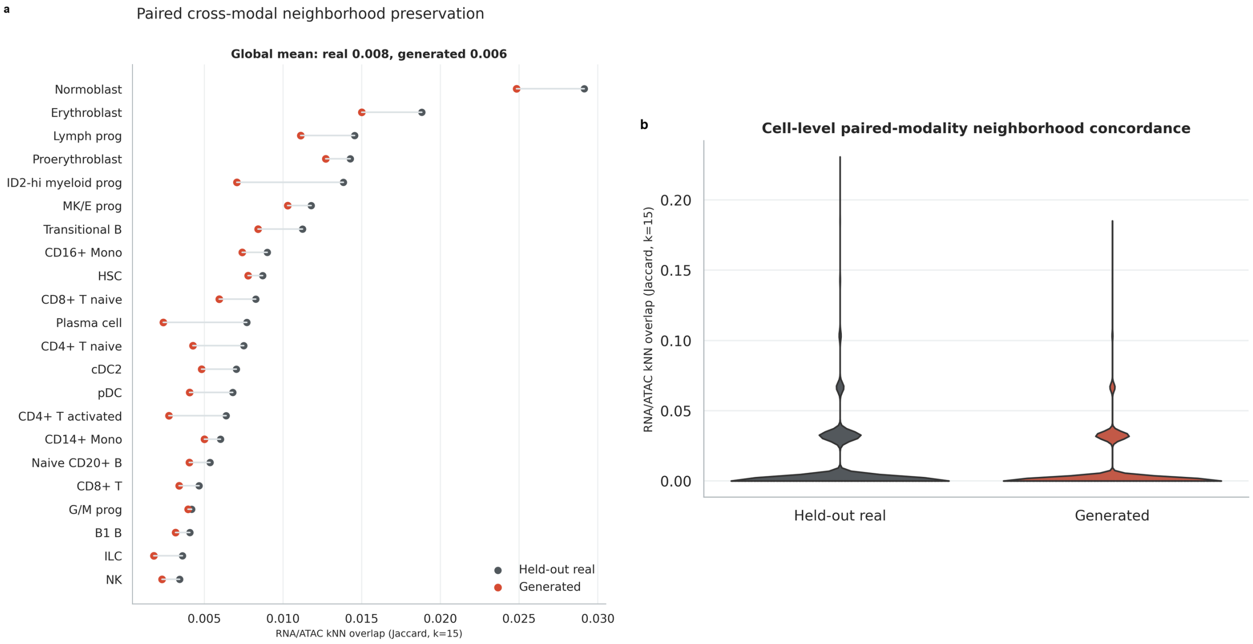


**Supplementary Figure 3**. Paired cross-modal neighborhood structure. a, Cell-type-resolved Jaccard overlap between RNA- and ATAC-defined nearest-neighbor sets in held-out real and generated cells. b, Cell-level distributions of paired-modality neighborhood concordance. These geometric analyses assess preservation of paired cross-modal organization and do not establish direct regulatory links.

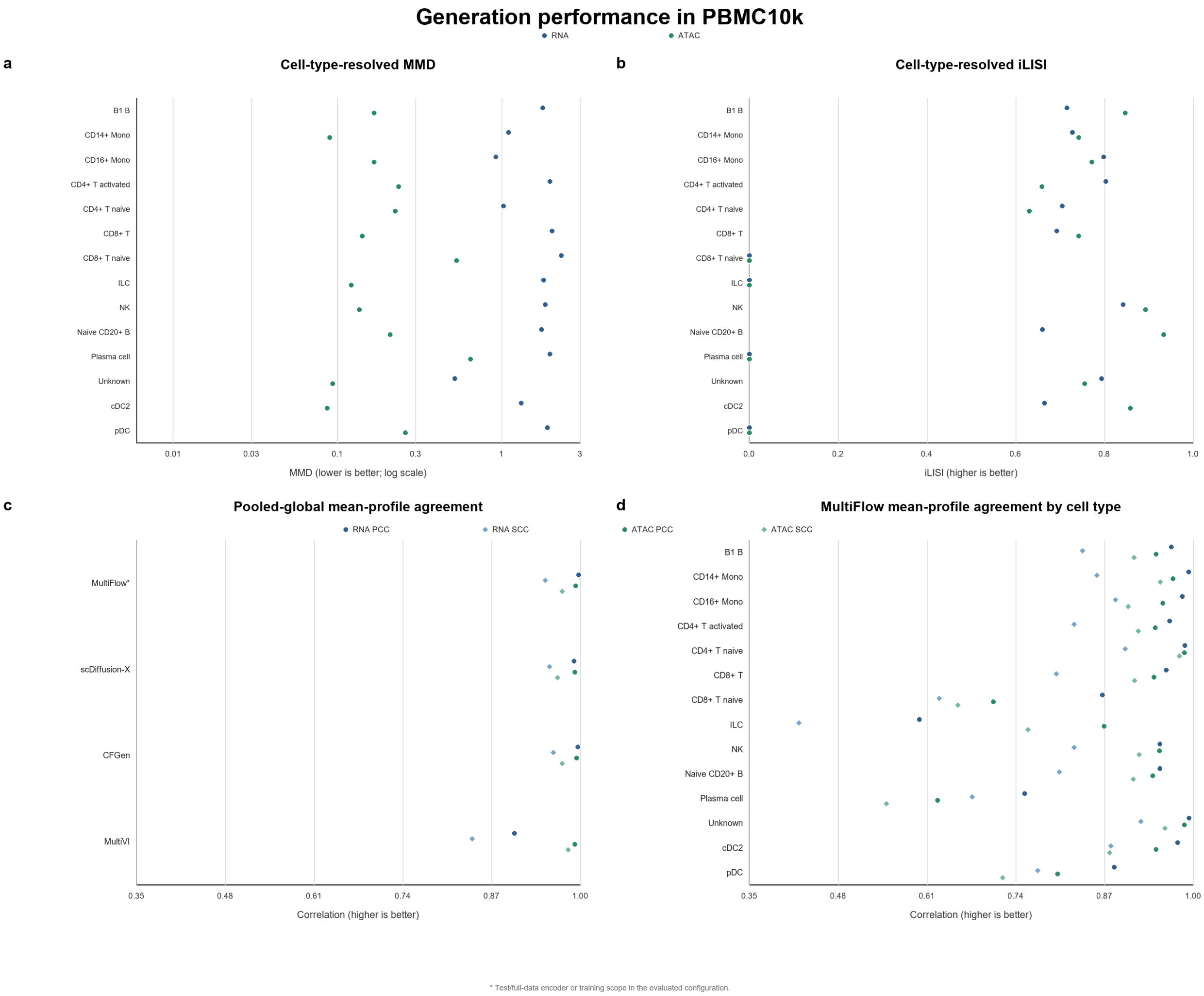


**Supplementary Figure 4. Generation performance in PBMC10k.** a and b, Cell-type-resolved maximum mean discrepancy (MMD) and integration local inverse Simpson's index (iLISI) for MultiFlow-generated RNA and ATAC profiles. Each point denotes one cell type. MMD is shown on a logarithmic scale; lower MMD and higher iLISI indicate better agreement between real and generated distributions. c, Pooled-global Pearson correlation coefficient (PCC) and Spearman correlation coefficient (SCC) across generation methods. d, Cell-type-resolved PCC and SCC for MultiFlow. Circles denote PCC, diamonds denote SCC, and colors distinguish RNA from ATAC; higher values indicate stronger agreement with real mean profiles.

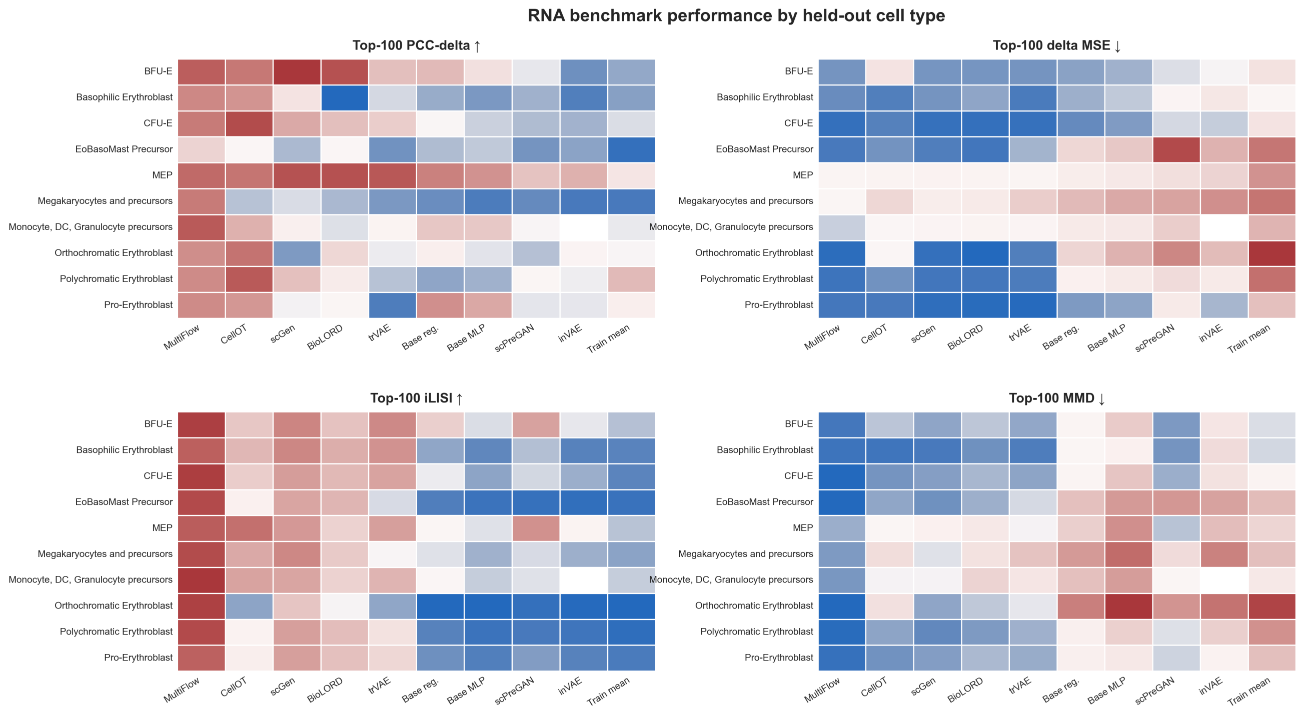


**Supplementary Figure 5.** RNA perturbation performance across held-out cell types. RNA benchmark performance across methods for the four primary endpoints reported in Fig. 3a: Top-100 PCC-delta, Top-100 delta MSE, Top-100 iLISI and Top-100 MMD. Heatmap colors are scaled independently within each metric; arrows indicate whether higher or lower values are better.

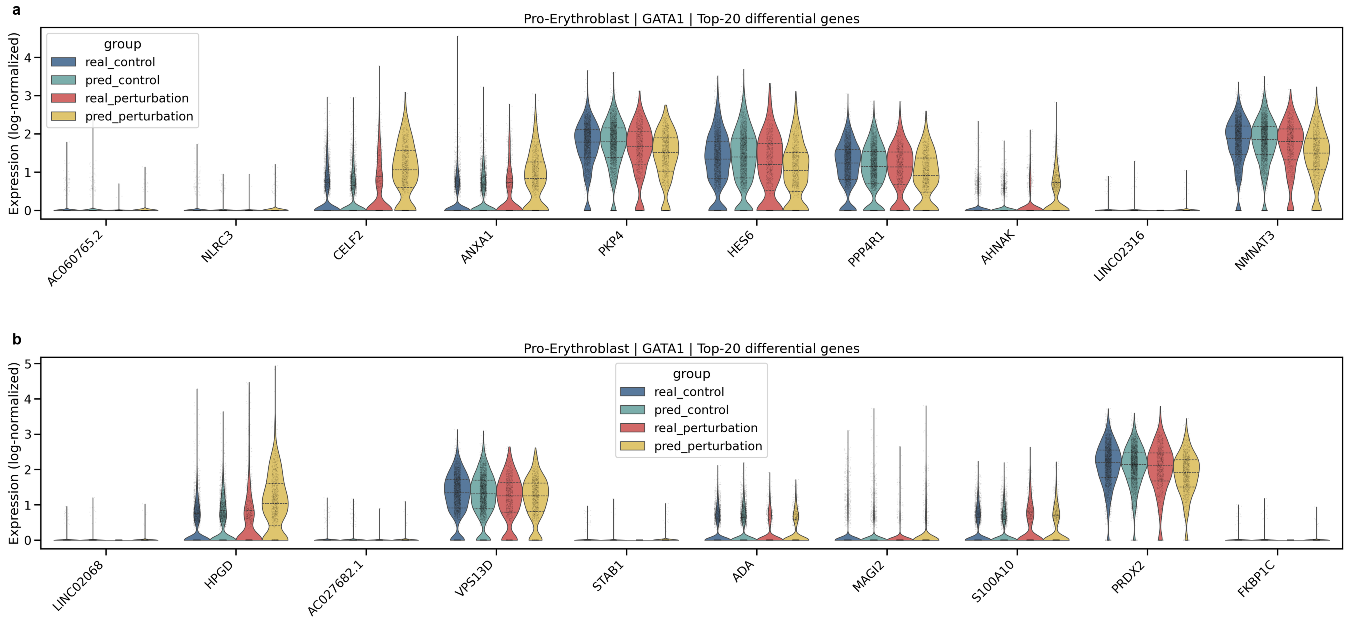


**Supplementary Figure 6.** Representative gene-level perturbation distributions. Expression distributions for the Top-20 GATA1-responsive genes in the held-out Pro-Erythroblast experiment. a,b, Real control, real perturbation, predicted control and predicted perturbation profiles are shown for the first and second groups of ten genes, respectively.

**All held-out cell types | ATF4 | RNA perturbation UMAP by cell type**

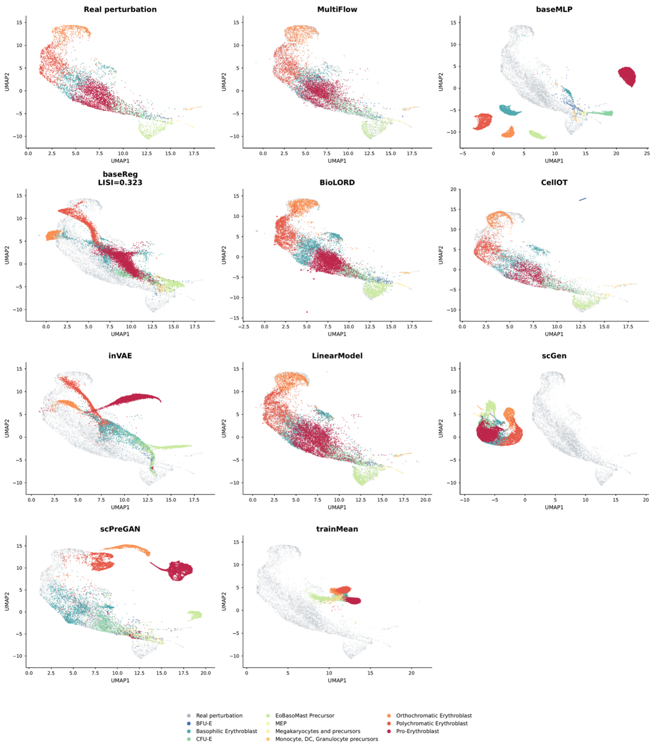


**All held-out cell types | BCL11A | RNA perturbation UMAP by cell type**

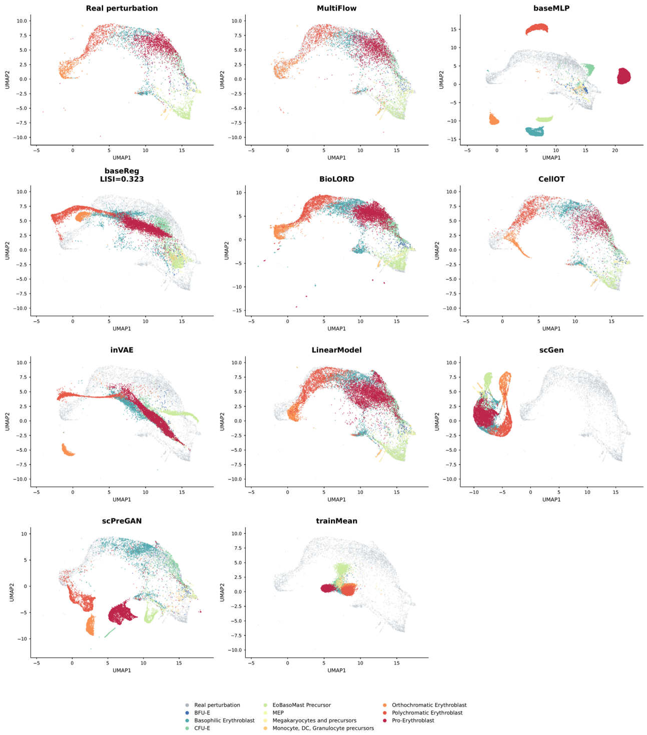


**All held-out cell types | FOSL1 | RNA perturbation UMAP by cell type**

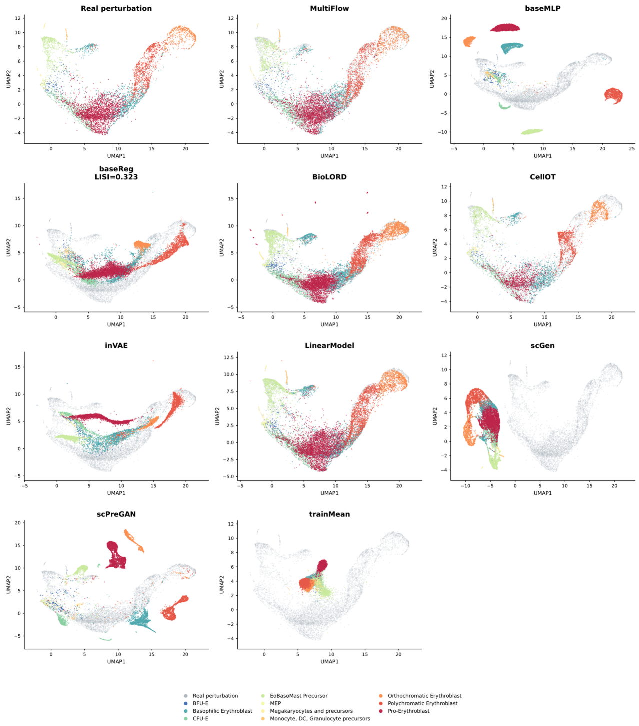


**All held-out cell types | GATA1 | RNA perturbation UMAP by cell type**

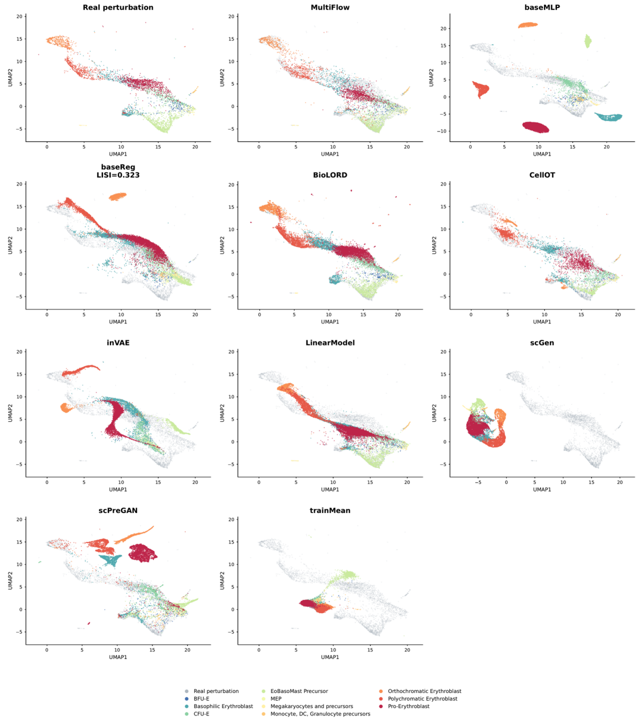

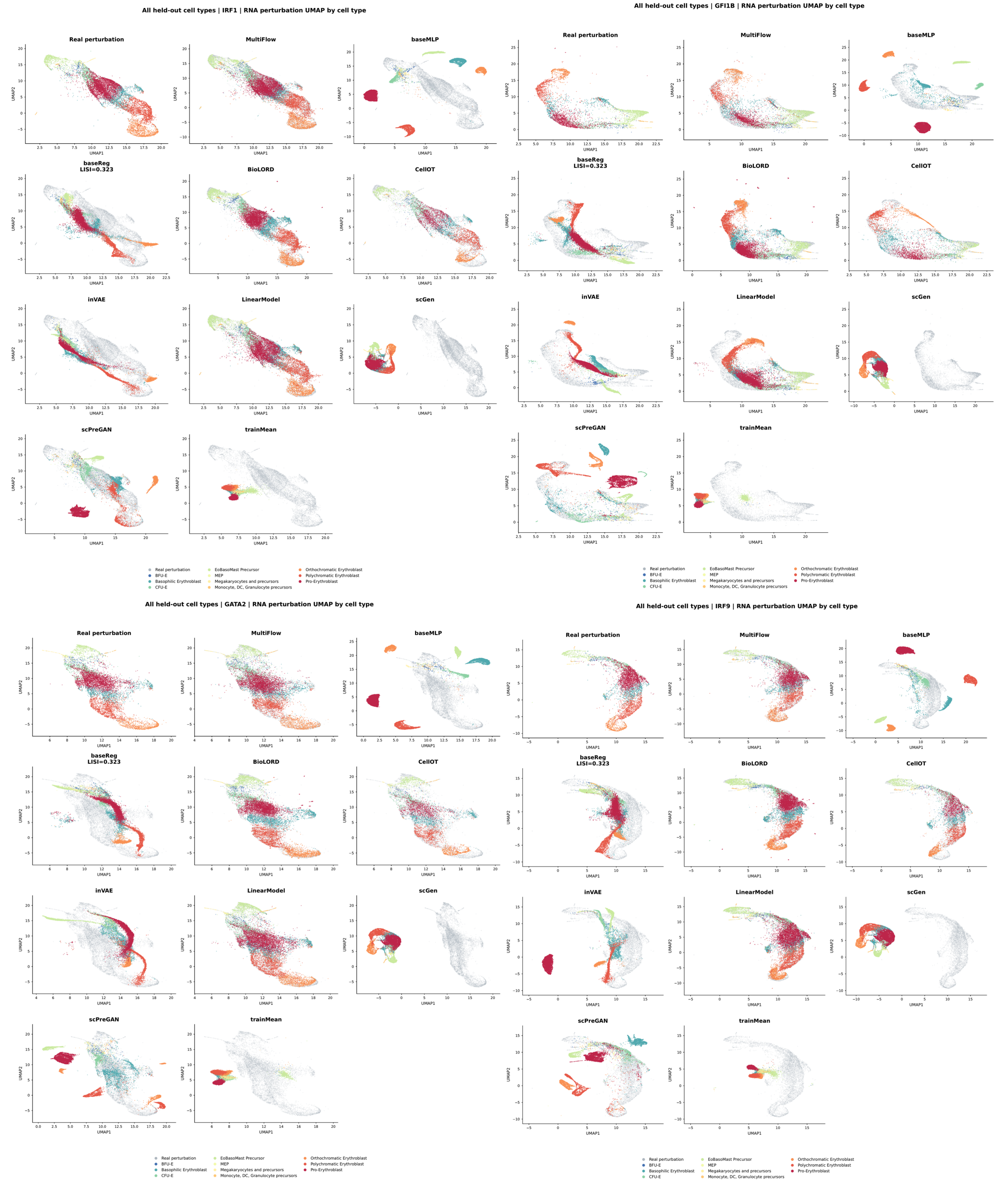


**Supplementary Figure 7.** RNA perturbation UMAP atlas. Shared-space RNA UMAPs for observed perturbed cells and all available prediction methods in the several benchmark. Cells are colored by held-out cell type. Real and predicted profiles were embedded under the same preprocessing and dimensionality-reduction protocol, enabling direct comparison of population coverage across methods.

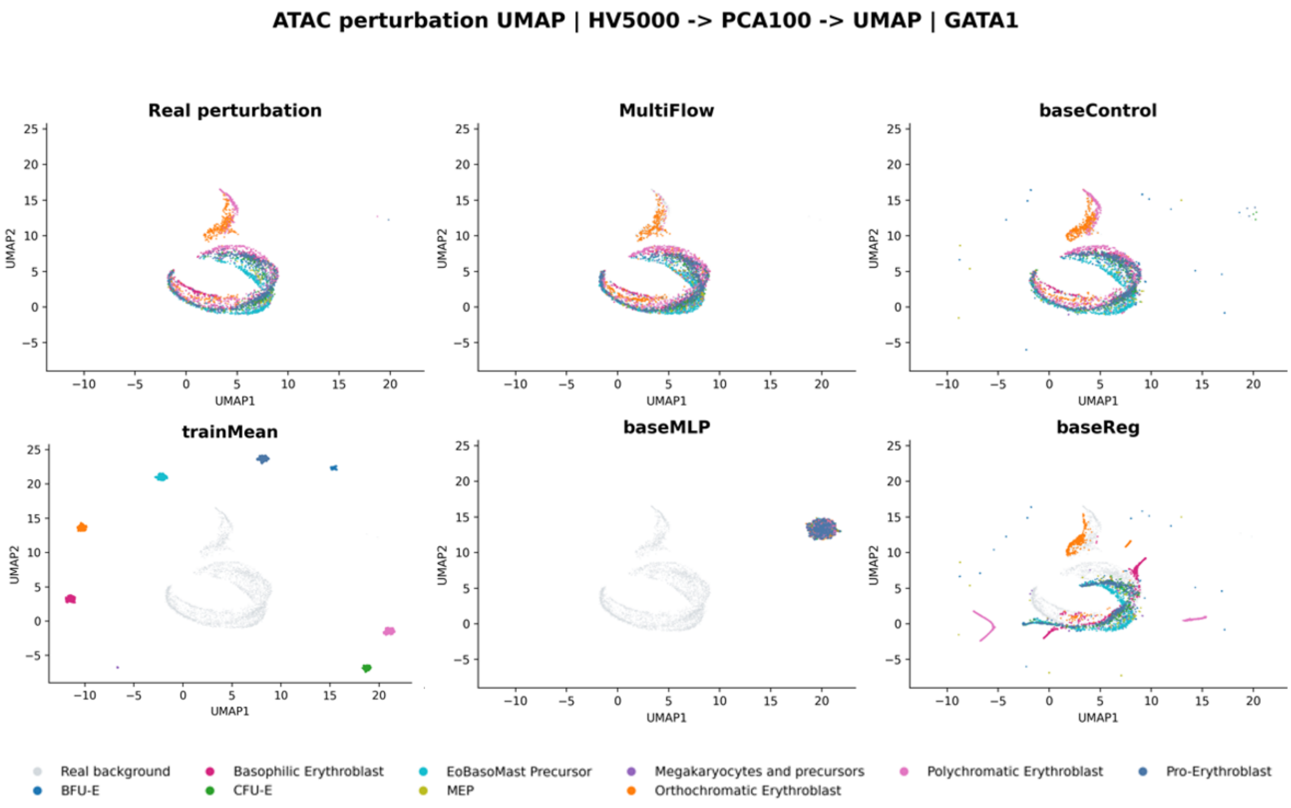


**Supplementary Figure 8.** ATAC perturbation UMAP atlas. Shared-space ATAC UMAPs for observed and predicted GATA1-perturbed cells using 5,000 highly variable peaks and 100 principal components. Panels show MultiFlow and the available paired multiomic baselines; cells are colored by held-out cell type.

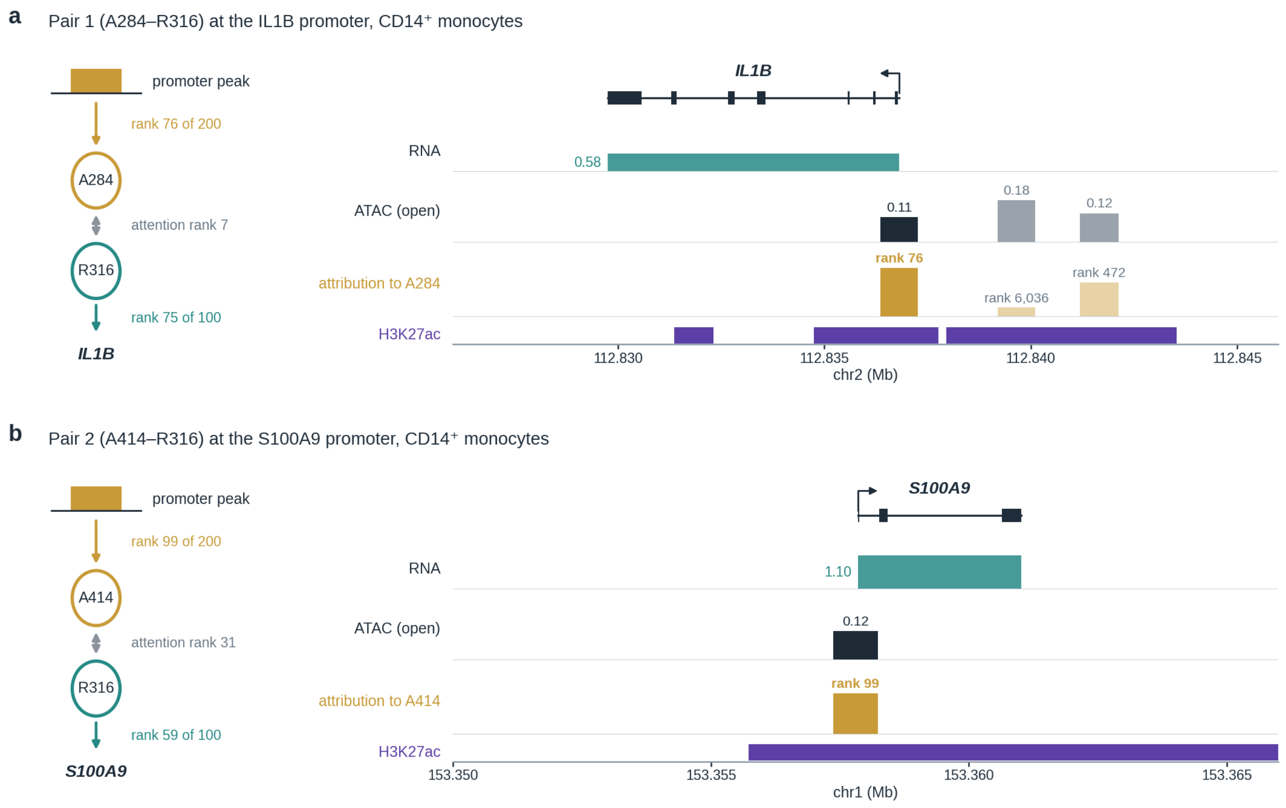


**Supplementary Figure 9.** The two example pairs of Fig. 4b at their loci in CD14$^{+}$ monocytes. a and b, Pair A284–R316 at the *IL1B* promoter (a) and pair A414–R316 at the *S100A9* promoter (b). Left, the model route: the promoter peak ranks 76th and 99th among the Top200 attributed peaks of the ATAC element, the pair ranks 7th and 31st among the 262,144 attention pairs, and the gene ranks 75th and 59th among the Top100 attributed genes of R316. In both loci the attributed peak spans the TSS and lies in H3K27ac; in the *IL1B* window it is the most strongly attributed of the three peaks.

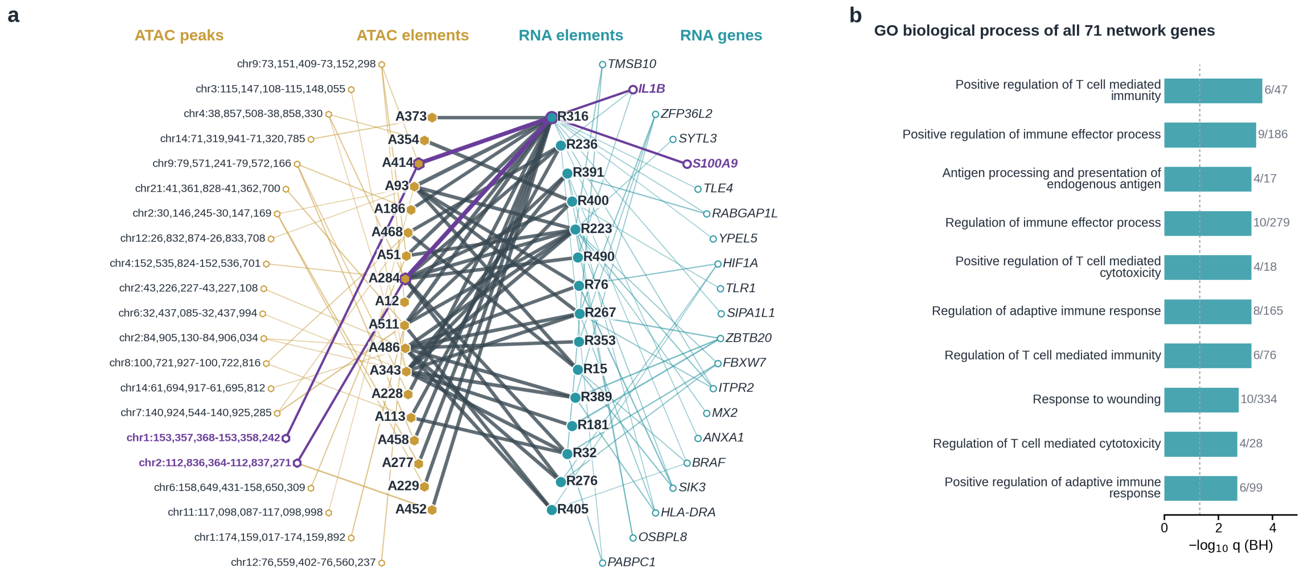


**Supplementary Figure 10.** A regulatory network of CD14$^+$ monocytes reconstructed from the attention map. a, Paths of the top 0.1% of attention pairs (263 of 262,144) in which a Top200 peak of the ATAC element lies at the promoter of a Top100 gene of the RNA element and overlaps CD14$^+$ monocyte H3K27ac: 245 paths from 152 pairs, reaching 71 genes. Shown are the 20 genes with the highest path score (attention × gene attribution × peak attribution), each with up to three paths; peaks are labelled by their hg38 coordinates. Gold, peak attribution; dark grey, attention between ATAC and RNA elements; teal, gene attribution; line width scales with attribution. Purple, pairs IL1B and S100A9 of Fig. 4b. b, The ten most significant of the 248 GO biological-process terms at $q < 0.05$ among the 71 genes (one-sided hypergeometric test against 13,431 model genes; BH).

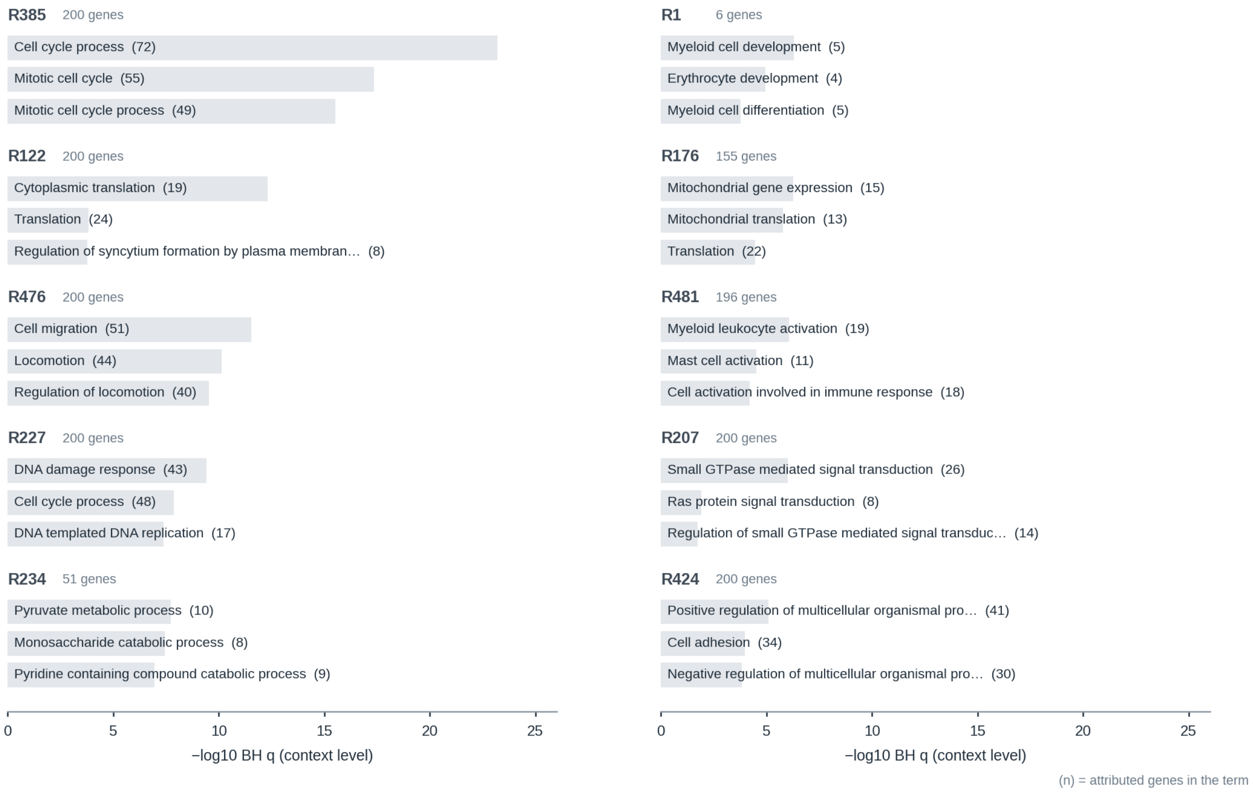


**Supplementary Figure 11.** Functional programs of individual RNA hidden elements (basophilic erythroblasts after GATA1 perturbation). Ten elements were chosen by a fixed rule: elements ranked by their best multi-hit GO term (context-level BH q), at most one element per gene-overlap cluster, and no element whose best term overlaps (gene Jaccard $\geq 0.25$) that of an element already kept. Bars, the three best terms (−log10 q; in brackets, attributed genes in the term). Programs include cell cycle (R385, 72 genes, q = 6e-24), cytoplasmic translation (R122), cell migration (R476), DNA damage response (R227), pyruvate metabolism (R234), myeloid and erythrocyte development (R1), mitochondrial gene expression (R176), myeloid leukocyte activation (R481) and small-GTPase signalling (R207).

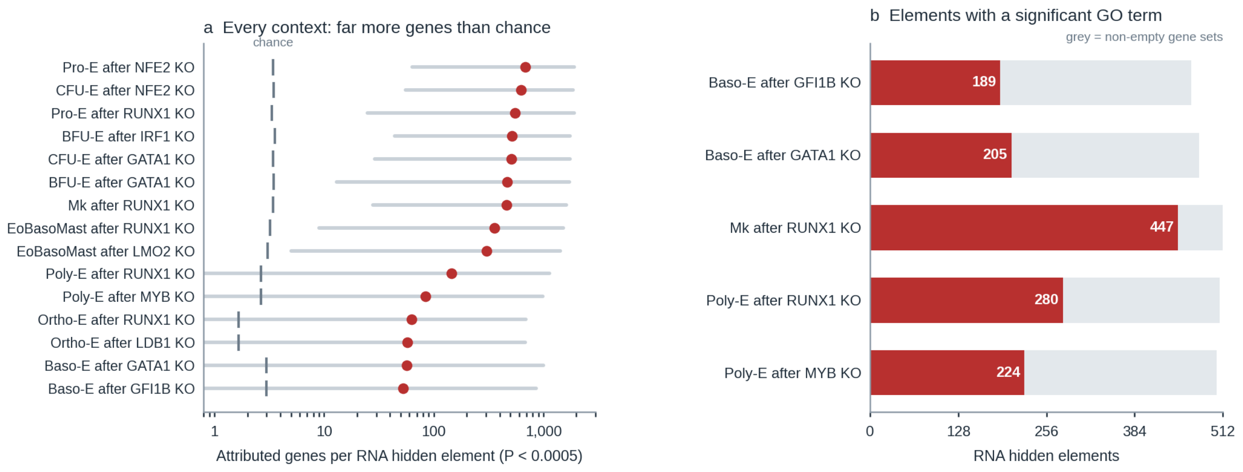


**Supplementary Figure 12.** Attributed gene sets and their GO signal across contexts. a, Genes per RNA hidden element at random-element $P < 0.0005$ in 15 training contexts: dot, median over the 512 elements; line, range; tick, genes expected by chance. b, Elements with at least one multi-hit GO term at context-level BH $q < 0.05$ in the five original contexts (189–447 of 512); grey, non-empty Top200 gene sets.

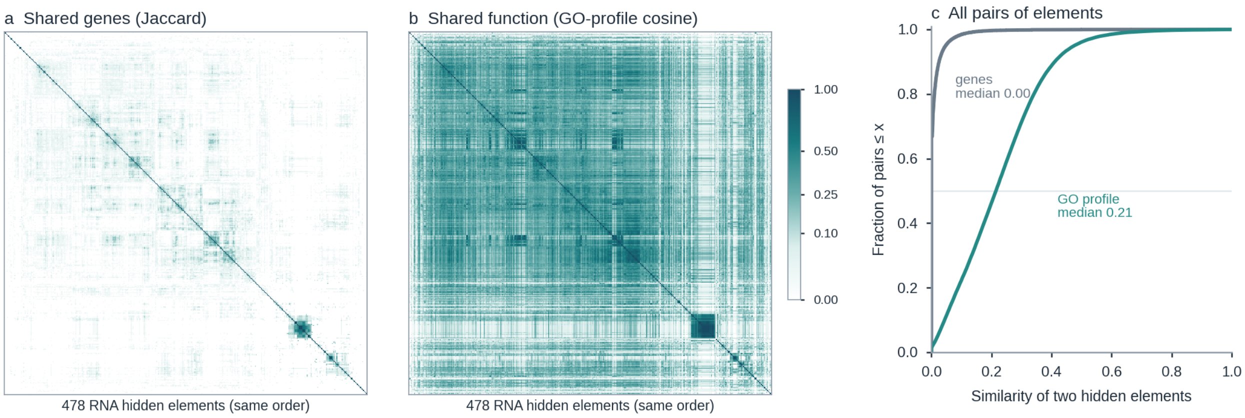


**Supplementary Figure 13.** RNA hidden elements share function more than genes (basophilic erythroblasts after GATA1 perturbation). a and b, Gene-set Jaccard similarity (a) and cosine similarity of GO profiles (b; −log10 raw P over all tested terms) for the 478 non-empty Top200 gene sets, in the same clustering order. c, Distribution of the off-diagonal values: median Jaccard 0.00, median GO-profile cosine 0.21.

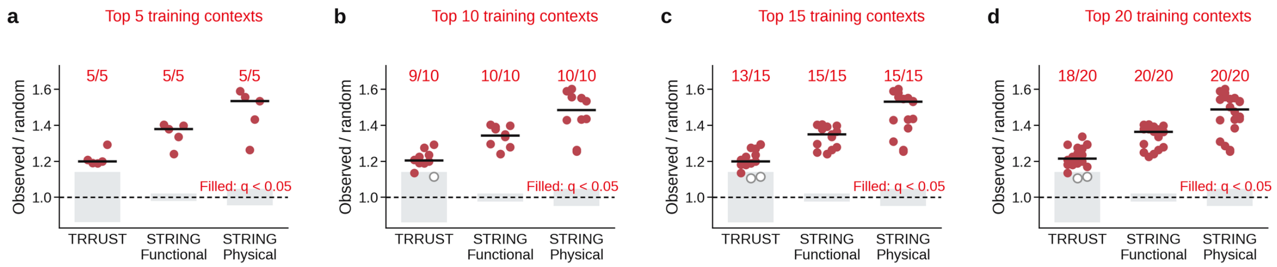


**Supplementary Figure 14.** Co-occurrence network enrichment does not depend on how many top-ranked training contexts are included. Nested sets of the top 5 (a), 10 (b), 15 (c) and 20 (d) sampling-eligible training contexts ranked by RNA PCC-delta; d is the set of Fig. 4h. Statistics as in Fig. 4h, with BH correction within each set; filled dots, $q < 0.05$. TRRUST is significant in 5/5, 9/10, 13/15 and 18/20 contexts, and STRING functional (≥ 400) and physical (≥ 700) in every context of every set (median fold, TRRUST / functional / physical: 1.20–1.22 / 1.34–1.38 / 1.48–1.53). The two non-significant TRRUST tests of the top 20 are orthochromatic erythroblasts after LDB1 perturbation ($P = 0.053$) and basophilic erythroblasts after MYB perturbation ($P = 0.078$).

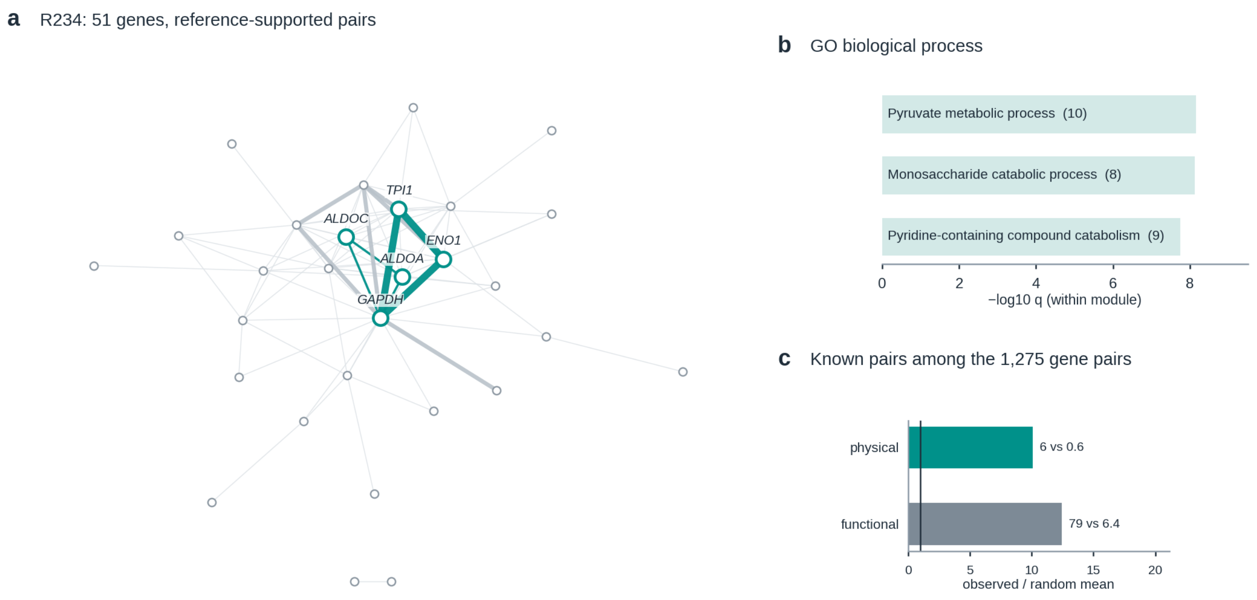


**Supplementary Figure 15.** a, Pairs among its 51 genes (teal, STRING physical ≥ 700; grey, STRING functional ≥ 400 only); thick edges, pairs co-selected by at least three hidden elements beyond gene frequency (hypergeometric $P < 0.01$). b, GO enrichment, BH within the module: pyruvate metabolic process (10 genes, $q = 7e\text{-}9$). c, 6 physical and 79 functional pairs among the 1,275 gene pairs, versus 0.6 and 6.4 expected by random paired (BH $q = 0.0048$ and $0.0006$).

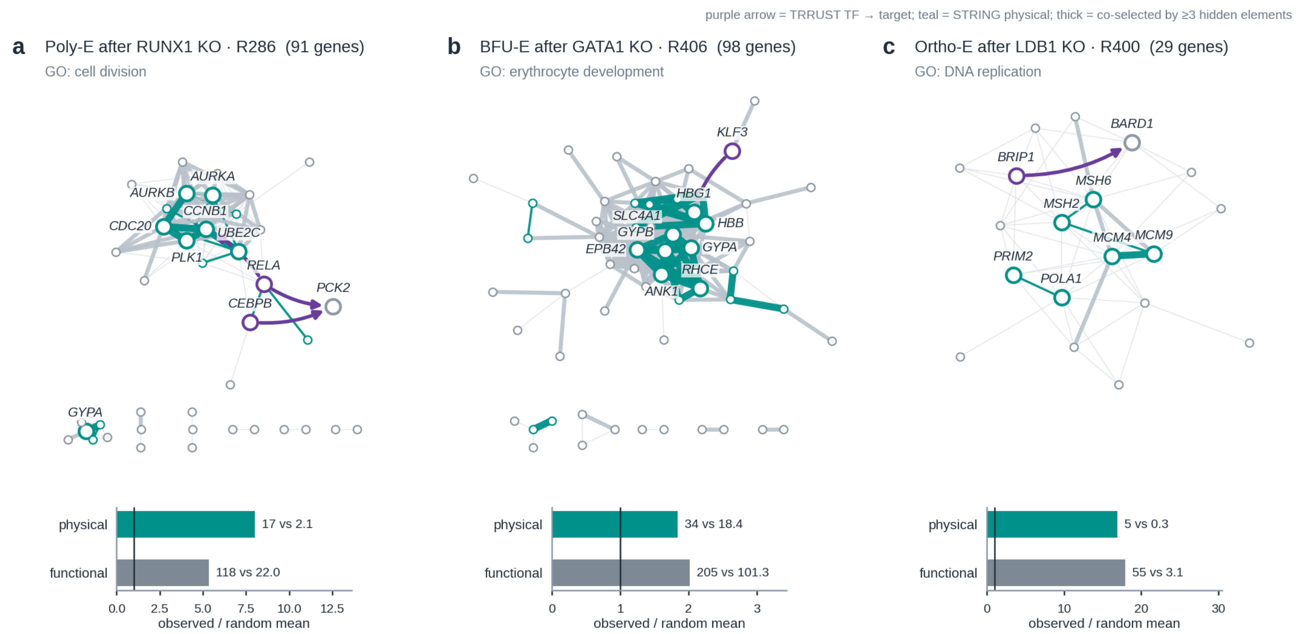


**Supplementary Figure 16.** Hidden-element gene modules supported by complementary reference databases. Three modules from different contexts: R286 of polychromatic erythroblasts after RUNX1 perturbation (cell division; 91 genes), R406 of BFU-E cells after GATA1 perturbation (erythrocyte development; 98 genes) and R400 of orthochromatic erythroblasts after LDB1 perturbation (DNA replication; 29 genes). Edges, co-selected gene pairs with reference support (purple arrows, TRRUST transcription factor → target; teal, STRING physical ≥ 700; grey, STRING functional ≥ 400 only); thick edges, pairs co-selected by at least three hidden elements more often than expected from gene frequency (hypergeometric $P < 0.01$). Bars, observed versus mean of 9,999 frequency-stratified permutations: physical and functional pairs 17 and 118 versus 2.1 and 22 (R286), 34 and 205 versus 18 and 101 (R406), and 5 and 55 versus 0.3 and 3.1 (R400).

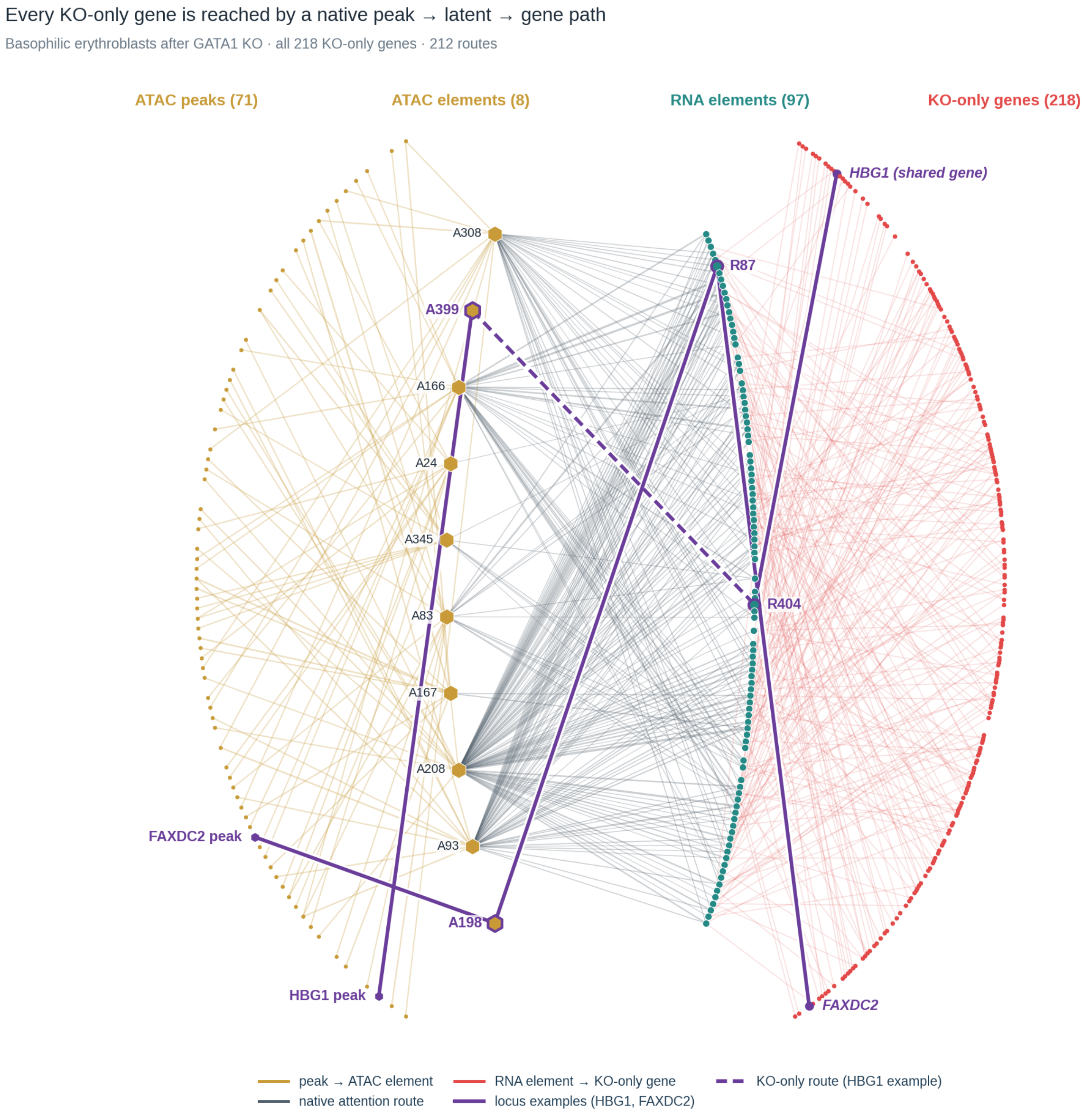


**Supplementary Figure 17.** Every knockout-only gene is reached by a native peak → latent → gene path (basophilic erythroblasts after GATA1 perturbation). The display rule of Fig. 5c applied to all 218 knockout-only genes: for each gene, its three highest-scoring routes and the Top10 peaks of each selected ATAC element, giving 212 routes through 8 ATAC and 97 RNA elements and 71 peaks. Purple, two locus examples added outside the rule: the HBG1 path through A399–R404, a route present only after knockout (dashed), and the FAXDC2 path through A198–R87.

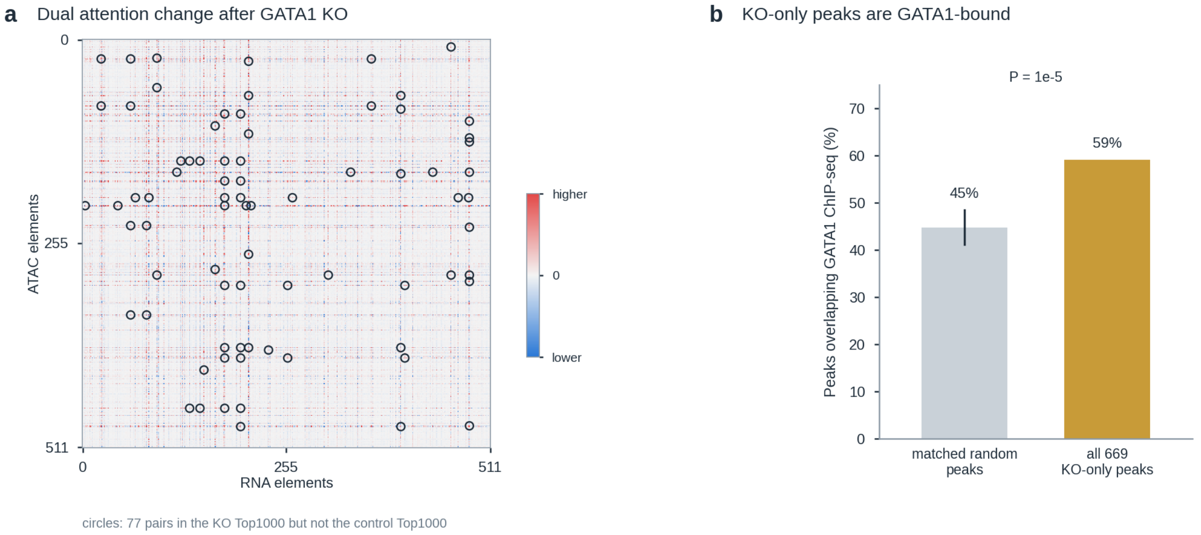


**Supplementary Figure 18.** Knockout-only attention pairs and their peaks (native network of basophilic erythroblasts after GATA1 perturbation). a, Dual attention after GATA1 knockout minus control; circles mark the 77 element pairs in the knockout Top1000 but not the control Top1000. b, 59% of the 669 knockout-only peaks overlap GATA1 ChIP-seq, versus 45% for 99,999 random peak sets matched for chromosome, accessibility and width (bar, central 95%; P = 1e-5).

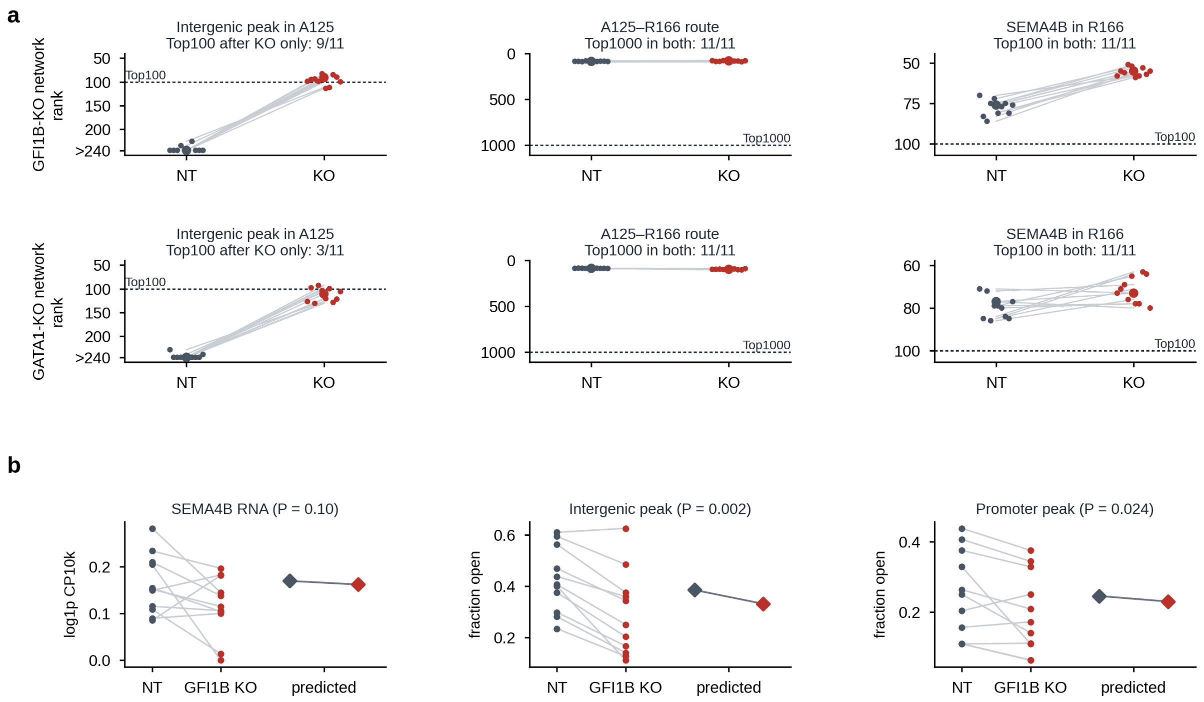


**Supplementary Figure 19.** The SEMA4B link of Fig. 5e. a, Ranks of the three steps of the link (intergenic peak in ATAC element A125, A125–R166 attention route, SEMA4B in RNA element R166) in NT and knockout (KO) cells, in the GFI1B-knockout network of Fig. 5e (top) and the GATA1-knockout network (bottom; purple in Fig. 5c). Large dots, full sample; small dots, ten half-samples; dashed lines, network cut-offs; peak ranks beyond 240 are drawn as >240. b, SEMA4B RNA and accessibility of the intergenic and promoter peaks per replicate stratum in NT and GFI1B-knockout cells (11 strata; two-sided Wilcoxon signed-rank test, descriptive), with the model prediction (diamonds). The link was selected from the 163 perturbation-only links with a GATA1 HiChIP contact to a transcription start site by a fixed rule: intergenic peak; loop in both replicates of a stage; perturbation-only in ≥ 6 of 11 samples; and real and predicted ATAC and RNA changes of the same sign.

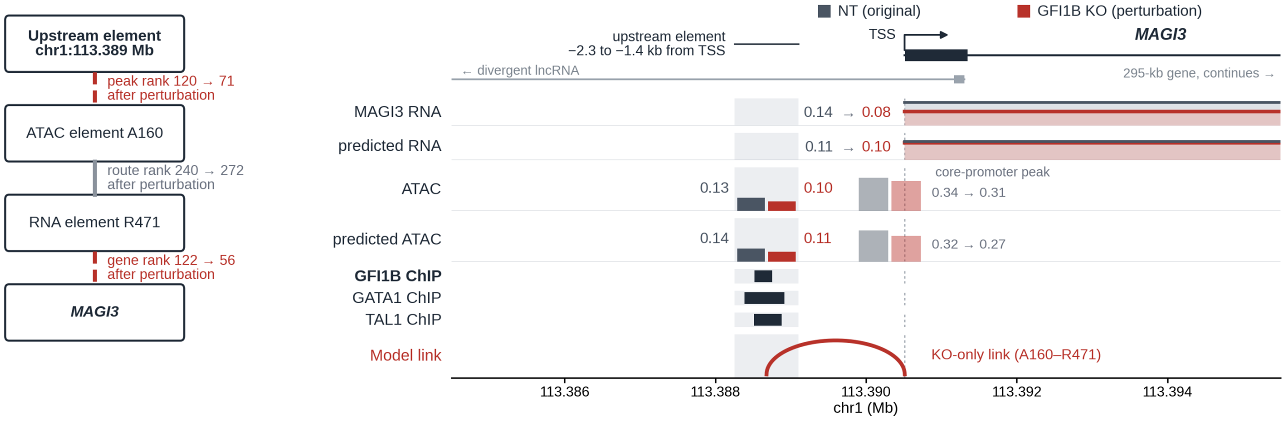


**Supplementary Figure 20.** A perturbation-only link from a factor-bound element upstream of the MAGI3 promoter (native network of basophilic erythroblasts after GFI1B perturbation). Left, the model route: the element (chr1:113,388,254–113,389,106; 1.4–2.3 kb upstream of the MAGI3 TSS) enters the Top100 peaks of ATAC element A160, and MAGI3 the Top100 genes of R471, only after GFI1B knockout, whereas the A160–R471 route is in the Top1000 of both states. Right, observed and predicted RNA and accessibility in NT (grey) and GFI1B-knockout (red) cells, ChIP-seq peaks and the model link.

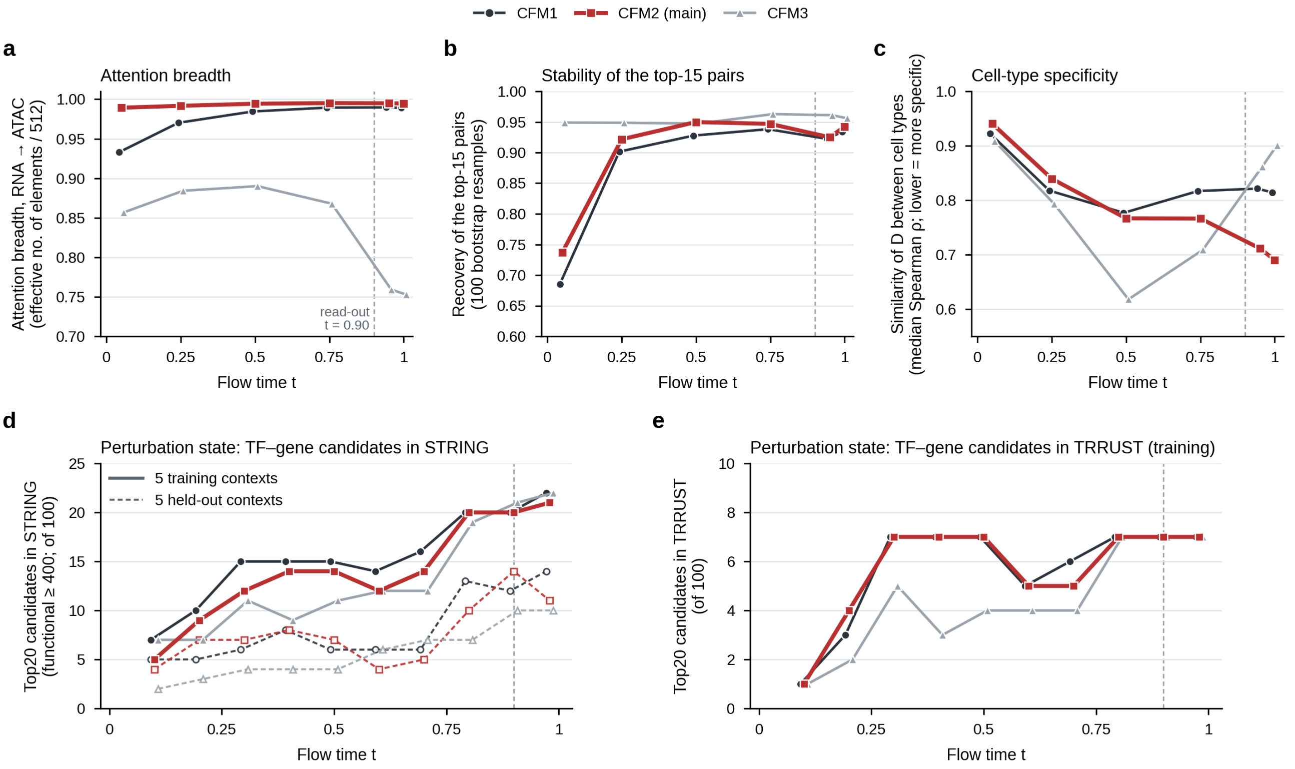


**f** Hub structure and marker recovery at other modules and times

| Module, time | Element shared by the 15 strongest pairs | ATAC importance breadth (1 = uniform) | Marker recovery (cell types, BH q < 0.05) |
|---|---|---|---|
| CFM1, t = 0.90 | R316, 10/15 | 1.00 | 21/22 † |
| CFM1, t = 1.0 | R316, 13/15 | 1.00 | – |
| **CFM2, t = 0.90** | R316, 8–14/15 ‡ | – | 22/22 |
| CFM2, t = 1.0 | R316, 15/15 | 1.00 | – |
| CFM3, t = 0.90 | A316, 15/15 | 0.66 | – |
| CFM3, t = 1.0 | A316, 15/15 | 0.60 | 21/22 † |

R, RNA element; A, ATAC element; columns 2 and 3, $CD4^+$ activated T cells. † Earlier protocol (Methods). ‡ Range over the 22 cell types.

**Supplementary Figure 21.** Effect of flow time and cross-flow module on the attention results. Figs 4 and 5 use the second cross-flow module (CFM2, red) at t = 0.90 (dashed line); the other two modules are shown for comparison. a–c figures use OpenProblems cells. a, How evenly RNA elements spread their attention over ATAC elements. b, How often the 15 strongest attention pairs are recovered when cells are resampled. c, Similarity of attention maps between cell types (lower, more cell-type-specific). d and e, Perturbation data: number of top-20 predicted targets of the perturbed TF that are recorded in STRING (d) or TRRUST (e); solid, training contexts; dashed, held-out contexts (d only). f, Hub element, spread of ATAC-element importance and marker-gene recovery (number of the 22 cell types) for each module and time. Details in Supplementary Note.

**Supplementary Table 1.** External reference data used in the interpretation analyses. a, Factor-binding references. One reference per perturbed factor was fixed before the overlap tests and used for all cell types (Fig. 5d); the GATA1 reference was also used in Supplementary Fig. 18, and the GATA1, TAL1 and GFI1B references for the ChIP tracks in Fig. 5e and Supplementary Fig. 20. b, Other reference data [1-3].

**a**, Factor-binding references

| Factor | Assay | Accession | Experiment | Cells | Peak set | Peaks | PMID |
|---|---|---|---|---|---|---|---|
| ATF4 | ChIP-seq | GSE143961 | GSE143961.ATF4.CD34-pos | Primary $CD34^+$-derived erythroblasts | ReMap 2022 | 2,605 | 32299090 |
| BCL11A | CUT&RUN | GSE104676 | GSE104676.BCL11A.CD34_Day9_30min | Primary $CD34^+$-derived erythroid cells, day 9 | ReMap 2022 | 32,769 | 29606353 |
| FOSL1 | ChIP-seq | ENCSR000BMV | ENCFF087MFG | K562 | ENCODE | 8,192 | ENCODE |
| GATA1 | ChIP-seq | ENCSR000EXP | ENCFF180BYN | Primary peripheral-blood-derived erythroblasts | ENCODE | 34,942 | ENCODE |
| GATA2 | ChIP-seq | GSE60792 | GSE60792.GATA2.CD34_DMSO | Primary $CD34^+$-derived erythroid progenitors, vehicle | ReMap 2022 | 8,083 | 27073918 |
| GFI1B | ChIP-seq | GSE52924 | GSE52924.GFI1B.CD34 | Primary adult proerythroblasts | ReMap 2022 | 3,581 | 24395799 |
| IRF1 | ChIP-seq | ENCSR809JSD | ENCFF093BBG | K562, CRISPR-tagged IRF1 | ENCODE | 1,993 | ENCODE |
| IRF9 | ChIP-seq | ENCSR926KTP | ENCFF863WCN | K562 expressing tagged IRF9 | ENCODE | 9,316 | ENCODE |
| KLF1 | ChIP-seq | GSE43625 | GSE43625.KLF1.erythroid_R3R4 | Primary nucleated erythroid cells (R3/R4) | ReMap 2022 | 34,995 | 23341446; 27141965 |
| LDB1 | ChIPmentation | GSE142227 | GSE142227.LDB1.K-562 | K562 | ReMap 2022 | 22,493 | 32960220 |
| LMO2 | ChIPmentation | GSE231422 | GSM7277440 | Primary bone-marrow MEPs | Author peaks | 24,312 | 37595278 |
| MYB | ChIP-seq | GSE90769 | GSE90769.MYB.THP-1 | THP-1, untreated | ReMap 2022 | 58,289 | 29590629 |
| NFE2 | ChIP-seq | GSE43625 | GSE43625.NFE2.erythroid_R3R4 | Primary nucleated erythroid cells (R3/R4) | ReMap 2022 | 28,478 | 23341446; 27141965 |
| PRDM16 | ChIP-seq | GSE289855 | GSM8800306 | NUP98::NSD1, WT1-knockout leukemia cells derived from fetal-liver HSPCs | Author peaks | 8,902 | 40166161; 41351880 |
| RUNX1 | ChIP-seq | GSE70660 | GSE70660.RUNX1.CD34_ADULT | Primary adult bone-marrow $CD34^+$ HSPCs | ReMap 2022 | 13,524 | 26766440 |
| SPI1 | ChIP-seq | GSE70660 | GSE70660.SPI1.CD34_FETAL | Primary fetal-liver $CD34^+$ HSPCs | ReMap 2022 | 65,013 | 26766440 |
| TAL1 | ChIP-seq | GSE52924 | GSE52924.TAL1.CD34 | Primary adult proerythroblasts | ReMap 2022 | 28,572 | 24395799 |

Peak set: ReMap 2022, uniformly processed hg38 peaks of the named experiment; ENCODE, released IDR-thresholded peaks (GRCh38); author peaks, hg38 peak calls deposited by the authors in GEO. Peaks, intervals on chr1–22, X and Y after removing duplicates. PMID, PubMed ID listed in the GEO series. MEP, megakaryocyte–erythroid progenitor; HSPC, hematopoietic stem and progenitor cell.

**b**, Other reference data

| Resource | Source and version | Details | Used in |
|---|---|---|---|
| H3K27ac ChIP-seq peaks | scDiffusion-X repository (Luo et al.), evaluation_script/encode; distributed there as ENCODE H3K27ac peaks, without experiment accessions | Six lineage-level files: CD14 ($CD14^+$ Mono), myeloid (ID2-hi myeloid prog), B cell (Naive $CD20^+$ B, B1 B, Transitional B), CD4 ($CD4^+$ T naive, $CD4^+$ T activated), CD8 ($CD8^+$ T, $CD8^+$ T naive), NK (NK) | Fig. 4e; Supplementary Figs 9 and 10 |
| GATA1 HiChIP | GEO GSE214807 (PMID 37254808): GSM6616188 and GSM6616189 (day 5), GSM6616190 and GSM6616191 (day 12) | Cord-blood $CD34^+$ cells differentiated ex vivo (day-5 erythroid progenitors, day-12 erythroid precursors), two replicates per day; author FitHiChIP loops at 5-kb resolution, lifted from hg19 to hg38 | Fig. 5e; Supplementary Fig. 19 |
| Gene annotation | GENCODE release 47 | Gene starts, promoters (3 kb upstream of the gene start) and transcript TSSs | Figs 4d and 5e; Supplementary Figs 9, 10, 19 and 20 |
| Gene Ontology | MSigDB C5 GO:BP, v2026.1 | Human biological-process gene sets with 15–500 tested genes | Fig. 4f,i; Supplementary Figs 10–13, 15 and 16 |
| TRRUST | v2, human (trrust_rawdata.human.tsv) | Transcription factor–target pairs | Fig. 4h,i; Supplementary Figs 14–16 and 21 |
| STRING | v12.0, Homo sapiens | Functional links (combined score ≥ 400) and physical links (score ≥ 700) | Fig. 4h,i; Supplementary Figs 14–16 and 21 |

### Supplementary Notes

#### Choice of cross-flow module and flow time for the attention analyses

MultiFlow couples the RNA and ATAC flows through three cross-flow modules (CFM1–CFM3), and each module produces cross-attention maps at every flow time $t$. The analyses of Figs. 4 and 5 read the cell-averaged attention map of CFM2 at $t = 0.90$ (Methods). This note explains the choice and describes how the three modules were compared across flow times, in cell-state data (OpenProblems) and in the perturbation state (Supplementary Fig. 21).

We used $t = 0.90$ rather than 1.0 because the midpoint integrator evaluates the velocity at $t = 0.90$ but never at $t = 1.0$. Late in the flow, the third CFM (CFM3) concentrated attention on a single ATAC element, whereas the first CFM (CFM1) and CFM2 gave similar results and CFM2 attention differed more between cell types. The module and flow time were fixed before the analyses of Figs. 4 and 5.

For Supplementary Fig. 21, attention maps of the three CFMs were computed for 32 OpenProblems cells of each cell type, with two fixed Gaussian sources, at $t = 0.05$, 0.25, 0.50, 0.75, 0.9 and 1.0. Attention breadth was the effective number of attended elements, exp(entropy), divided by 512; stability was the mean frequency with which the 15 largest entries of $D$ were recovered in 100 bootstrap resamples of cells; and cell-type specificity was the median Spearman correlation of $D$ between cell types. ATAC-element importance was the mean attention that RNA elements pay to each ATAC element, and its breadth was computed in the same way ($CD4^+$ activated T cells). In the perturbation state, attention (192 generated samples per context) and output attribution (24 samples) were computed at $t = 0.1$–0.9 and 0.98 along saved generation trajectories. Target genes were ranked by attention-weighted output attribution over the 1,000 strongest pairs, and pairs of the perturbed factor with its 20 top-ranked genes that are recorded in STRING v12 (functional score $\geq 400$) or TRRUST v2 were counted over five training contexts and five held-out contexts whose perturbation response was withheld from training of the checkpoint.

In summary, late in the flow CFM2 spread its attention almost evenly over the ATAC elements (breadth 0.995), whereas CFM3 concentrated it on one element (breadth 0.75–0.87 for $t \geq 0.75$; Supplementary Fig. 21a). The 15 strongest CFM2 pairs were stable from $t = 0.25$ onward (recovery 0.92–0.95; Supplementary Fig. 21b), and at $t = 0.95$ CFM2 attention was the most cell-type-specific of the three modules (median Spearman ρ between cell types: CFM2, 0.71; CFM1, 0.82; CFM3, 0.86; Supplementary Fig. 21c). In the perturbation state, database support of the top-ranked targets increased with flow time and was similar for the three modules in training contexts at $t \geq 0.8$ (19–22 of 100 candidates in STRING and 7 of 100 in TRRUST; Supplementary Fig. 21d,e). Figs. 4 and 5 report CFM2 at $t = 0.90$ only; the other modules and flow times are shown for comparison.
